# Bovine AAV - a promising vector for pulmonary gene therapy

**DOI:** 10.64898/2026.07.14.738418

**Authors:** Daniela C. Ivan, Valerie Dubost, Laura Israel, Jonas Weinmann, Dennis Ungan, Nina Carbonetti, Nathalie Stuber, Magali Jivkov, Esther Erard, Elisabetta Biglieri, Federica De Girardi, Simon Mittermeier, Maryam Syed, Bruno Tigani, Najwa Ouali-Alami, Kristin Dreessen, Colin Deniston, Kannan Sankar, Laura Bollepalli, Vanessa Cornacchione, Elisabetta Traggiai, Dominique Brees, Anette Karle, José M. Carballido, Agostino Cirillo

**Author notes:** Correspondence should be addressed to: Daniela Condeescu-Ivan and Agostino, Cirillo. These authors have contributed equally to this work (José M. Carballido, Agostino Cirillo).

## Abstract

Efficient systemic delivery to the lung remains a major barrier for adeno-associated virus (AAV)-mediated pulmonary gene therapy, particularly when pre-existing immunity limits the use of conventional capsids. Here, we evaluated Bovine AAV, a phylogenetically divergent capsid, as candidate vector for lung-directed gene transfer. In adult C57BL/6J mice, intravenous delivery of Bovine AAV resulted in robust and preferential lung transduction comparable to AAV4, with predominant targeting of alveolar type I pneumocytes and pulmonary endothelial cells. In primary human lung-resident cells, Bovine AAV was particularly effective in microvascular endothelial cells, a target poorly transduced by AAV4 *in vitro*. Bovine AAV demonstrated scalable production with yield, purification performance, capsid quality, and genome integrity comparable to AAV9. In sera from healthy adults from the United States and Switzerland, Bovine AAV showed intermediate neutralization frequencies, lower than AAV2 and AAV4 but higher than AAV5 and AAV9. Of relevance, Bovine AAV maintained *in vivo* transduction efficiency in mice previously immunized with a pool of human and non-human primate-derived AAV capsids, including AAV4. Together, these results position Bovine AAV as a promising lung-tropic and immune-distinct vector for pulmonary gene therapy, with particular relevance for applications requiring systemic delivery in the presence of pre-existing immunity to conventional serotypes.

## Introduction

Adeno-associated virus (AAV)-mediated gene transfer has become a leading platform for *in vivo* gene therapy, yet efficient and durable delivery to the lung remains a major challenge and an unmet medical need. The pulmonary system is an attractive therapeutic target for a broad range of inherited and acquired diseases ^1^, but its intricate anatomy, extensive epithelial barriers, mucus and surfactant layers, and marked cellular diversity complicate vector access and restrict transduction efficiency (reviewed in ^2,3^). These constraints are compounded by the need to reach specific therapeutically relevant cell populations within the lung, a requirement that many currently available AAV capsids do not fulfill adequately ^4,5^. As a result, and despite substantial progress in capsid engineering strategies, there remains a need for AAV vectors that can efficiently access therapeutically relevant pulmonary cell populations while retaining properties compatible with clinical development ^1,4,5^.

A further obstacle to successful AAV-mediated gene therapy is pre-existing humoral immunity. Neutralizing antibodies against naturally circulating human AAV serotypes are widespread in the general human population ^6^ and can substantially reduce transduction efficacy ^7,8^, particularly after systemic administration ^9,10^. Beyond excluding a substantial fraction of potential patients ^11,12^, such immunity may restrict the use of related capsids with overlapping antigenic profiles ^9,10^, particularly in situations when gene therapy re-administration through a different vector is needed because of rapid cell turnover ^13^. Since many of the AAV vectors currently prioritized for translational development originate from human or non-human primate (NHP) isolates ^4,12^, they occupy a relatively narrow antigenic and evolutionary landscape, which can increase the likelihood of serological cross-reactivity^9^. For lung-directed gene therapy, where systemic delivery may be advantageous for reaching diffuse or vascular compartments, these immune constraints represent a particularly important obstacle. Therefore, identifying vectors outside this space may offer an opportunity to expand the repertoire of clinically relevant capsids with differentiated immune properties.

Evolutionary distant AAVs are especially interesting, as they may combine distinct tissue tropism with reduced serological overlap not captured by conventional human and NHP-derived serotypes. Bovine AAV represents one such candidate. This capsid is phylogenetically distinct from most widely studied AAVs ^14^ and has previously been reported to display lung-associated tropism ^15,16^. It shares substantial capsid similarity with AAV4 ^14^, one of the most structurally divergent serotypes derived from NHPs ^17^, but remains sufficiently distinct to warrant independent functional and immunological evaluation. This relationship is noteworthy because AAV4 has emerged as a lung-relevant benchmark in recent comparative studies ^18^, yet its overall translational profile remains incompletely resolved. Bovine AAV therefore raises the possibility that a naturally divergent capsid may provide a favorable combination of pulmonary tropism and serological differentiation.

Importantly, tropism alone is insufficient to establish translational value. A candidate vector for pulmonary gene therapy must be evaluated within a broader framework that includes *in vivo* performance, relevance across human target cell types, susceptibility to antibody-mediated cross-neutralization, prevalence of pre-existing immunity in human populations, and compatibility with scalable manufacturing processes ^4,9,19^. These considerations are especially important when evaluating unconventional capsids, where promising biological observations do not always translate into development feasibility. A rigorous assessment of Bovine AAV should therefore determine not only whether it can transduce the lung, but whether it offers a meaningful overall advantage relative to more established and emerging AAV comparators.

In this study, we performed a comprehensive preclinical evaluation of Bovine AAV as a candidate vector for lung-directed gene transfer. We investigated its *in vivo* transduction profile in mice after systemic administration and compared its performance with established AAV capsids, with particular emphasis on AAV4 as a phylogenetically related comparator. We then examined Bovine AAV from two complementary immunological perspectives: first, its susceptibility to cross-neutralization by antibodies generated *in vivo* against a panel of commonly used human and NHP-derived AAVs; and second, the prevalence of Bovine AAV-directed neutralizing antibodies in healthy adult human cohorts. To strengthen the translational relevance of this analysis, we evaluated Bovine AAV activity in primary human lung cells and examined its manufacturability and capsid quality at larger production scale. This study was designed to determine whether Bovine AAV represents a lung-tropic, serologically distinct and translationally viable alternative to conventional AAV platforms for pulmonary gene therapy.

## Results

### Bovine AAV drives robust and preferential lung transduction *in vivo*

Systemic delivery of AAV vectors is an attractive approach for pulmonary gene therapy, as it is minimally invasive, promotes homogeneous vector distribution, and facilitates access to distal lung regions, often difficult to reach via intranasal or intratracheal routes ^20^. Among natural serotypes, AAV4 shows the strongest lung transduction after intravenous administration in adult mice ^21^ and a robust lung transduction in crab-eating macaques ^18^. Among engineered capsids, AAV2-ESGHGYF stands out, a vector generated through *in vivo* peptide-display library selection, displaying enhanced and selective lung tropism in FVB/N mice following intravenous delivery ^22^. The phylogenetically distant Bovine AAV (Supplemental Figure 1, Supplemental Table 1), has also been reported to transduce lung tissue following intranasal administration, although this may depend on modulation of viral transcytosis and transduction may be transient ^15^. Notably, its tropism and transduction efficiency after systemic administration remains unknown to date.

Here, we sought to directly compare the pulmonary transduction efficiency of AAV4, AAV2-ESGHGYF, and Bovine AAV in C57BL/6J mice following intravenous delivery, using AAV9 as a comparator with limited lung tropism. Male mice aged 9-11 weeks received 1 × 10^13^vg/kg of each vector, corresponding to 2.5 × 10^11^vg per 25g mouse, packaged with an ss-CAG-GFP expression cassette (Figure 1a). Animals were monitored throughout the 4-week study, and no significant changes in body weight were observed in any vector-treated group relative to controls, indicating good tolerability under the conditions tested (Figure 1b).

**Figure 1.**
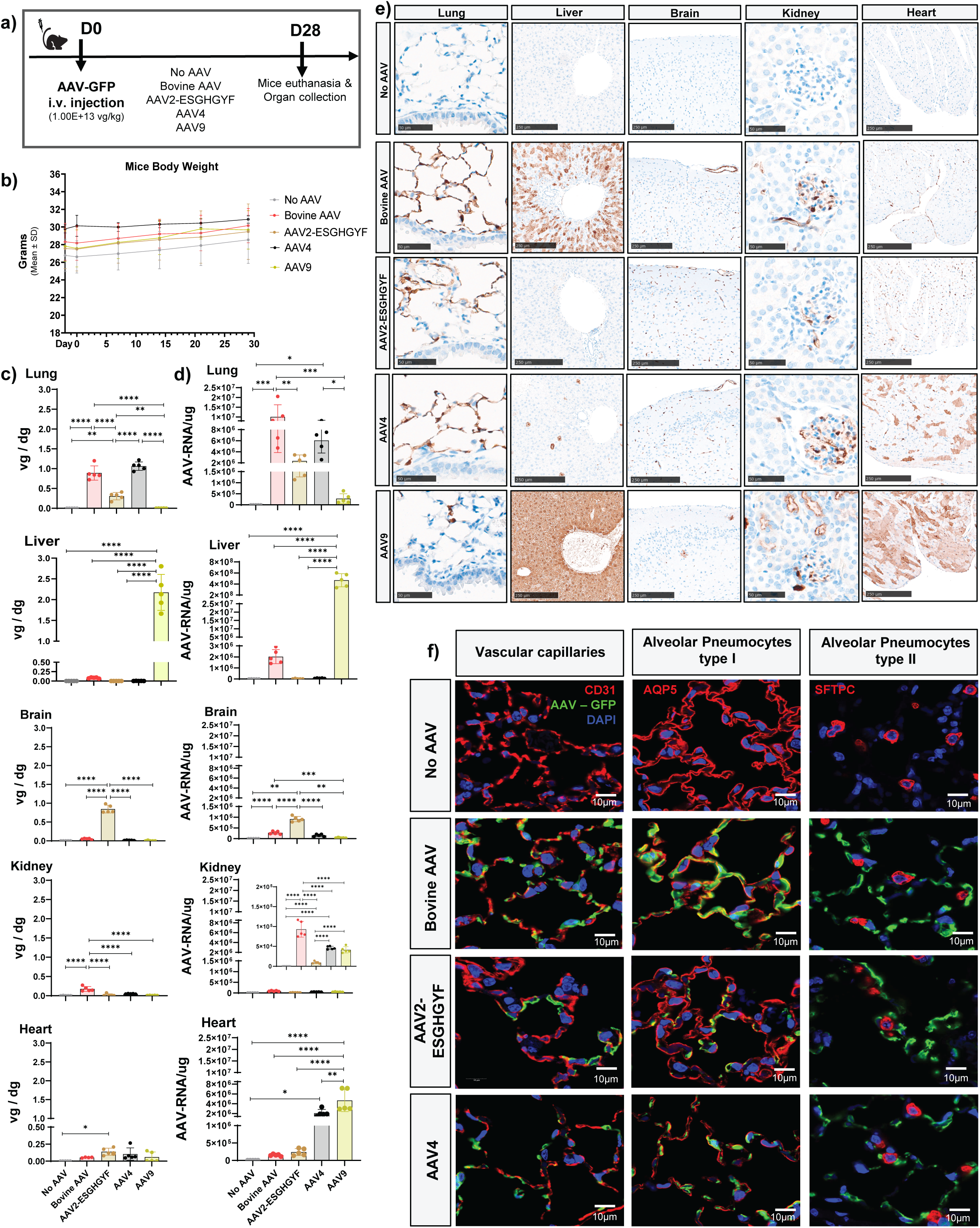
***In vivo* transduction efficacy of AAV2-ESGHGYF, AAV4, AAV9 and Bovine AAV. a) Study design.** Male C57BL/6J mice aged 9 to 11 weeks were injected intravenously via the tail vein with AAV2-ESGHGYF, AAV4, AAV9 or Bovine AAV vectors carrying an ss CAG GFP cassette at a dose of 1 × 10^13^ vg/ kg, corresponding to 2.5 × 10^11^ vg per 25 g mouse (*n* = 5 mice /group). An additional group of mice injected with AAV formulation buffer only (*n* = 5) served as negative controls. Four weeks after injection, mice were euthanized and the lung, liver, brain, kidney, and heart were collected for downstream analyses. **b) Body weight after AAV administration.** Body weight was recorded every 7 days throughout the study, with an additional baseline measurement obtained 3 days before vector administration. Data are shown as mean (SD) body weight in grams for n=5 mice per group. Differences in body weight changes over time depending on the injected AAVs were analyzed using a two-way repeated-measures ANOVA with AAV treatment group as the between-subject factor and time as the within-subject factor, followed by Sidak multiple-comparisons test. **c) Tissue vector biodistribution** and **d) Transgene expression.** c) Recombinant AAV vector genomes were quantified by ddPCR and are presented as vector genomes per diploid genome (vg/ dg), whereas d) Transgene expression was quantified by RT-ddPCR and is presented as AAV RNA per microgram of tissue. Because expression levels in the kidney were lower than in other organs, an enlarged view of the kidney data is included to facilitate comparison among serotypes. For panels **c** and **d**, points represent individual mice, and bars indicate mean (SD). Statistical significance was assessed by one-way ANOVA with Tukey‘s multiple comparisons test (∗p<0.05, ∗∗p<0.01, ∗∗∗p<0.001, ∗∗∗∗p<0.0001) **e) Representative histology images.** Representative immunohistochemistry images of formalin-fixed paraffin-embedded (FFPE) sections from lung, liver, brain, kidney, and heart illustrate localized cellular transduction following administration of Bovine AAV, AAV2-ESGHGYF, AAV4, or AAV9. The lung epithelial layer is emphasized with the dashed box. Scale bars, 50 μm for lung and kidney and 250 μm for liver, brain, and heart. **f) Co-localization of AAV2-ESGHGYF, AAV4 and Bovine AAV transduction with lung cell-type markers.** Double immunofluorescence staining and confocal microscopy were used to determine the cellular identity of transduced cells in the mouse lung following administration of Bovine AAV, AAV4, or AAV2 ESGHGYF. Lung tissue from mice receiving formulation buffer only (No AAV) was included as a negative control. GFP immunostaining (green) was used to identify vector transduced cells and was combined with staining for CD31, aquaporin 5 (AQP5) and surfactant protein C (SFTPC) (red) to identify endothelial cells, alveolar type I cells and alveolar type II cells, respectively. Nuclei were counterstained with DAPI (blue). Representative confocal images are shown. Scale bars, 10 μm.

Twenty-eight days after intravenous AAV injection, lung, liver, brain, kidney, and heart were collected for analysis. Vector biodistribution was quantified by droplet digital polymerase chain reaction (ddPCR) as vector genomes per diploid genome (vg/dg), transgene expression was assessed by reverse transcription ddPCR (RT-ddPCR) as viral RNA per microgram of tissue (AAV-RNA/μg) and GFP protein expression was evaluated by immunohistochemistry (IHC) on formalin-fixed, paraffin-embedded (FFPE) tissue sections (Figure 1c-e).

Bovine AAV and AAV4 showed the highest levels of lung transduction across the vectors tested (Figure 1c-e). At the RNA level, Bovine AAV yielded slightly higher expression than AAV4 although this difference was not statistically significant (Figure 1d). At the protein level, both capsids produced similar GFP staining patterns in the lung (Figure 1e). In contrast, AAV2-ESGHGYF showed substantially lower lung transduction than either Bovine AAV or AAV4, consistently across vector genome, RNA, and protein readouts (Figure 1c-e). Based on the tissue anatomy, we could already observe that none of the AAVs could penetrate and transduce the airway epithelial layer (Figure 1e). To define the cellular identity of transduced lung cells, FFPE lung sections were analyzed by double immunofluorescence staining and confocal microscopy using GFP to identify vector-transduced cells together with Aquaporin 5 (AQP5), Surfactant Protein C (SFTPC), and CD31 as markers of alveolar type I pneumocytes, alveolar type II pneumocytes, and endothelial cells, respectively. None of the AAV capsids tested showed detectable GFP co-localization with SFTPC, indicating no detectable transduction of type II pneumocytes (Figure 1f). Bovine AAV showed robust GFP co-localization with AQP5-positive alveolar type I pneumocytes and was also abundantly detected in CD31-positive endothelial cells (Figure 1f). Both AAV4 and AAV2-ESGHGYF were likewise detected in AQP5-positive cells, but less frequently (in fewer cells) than Bovine AAV. AAV4 also showed some co-localization with CD31-positive endothelial cells, whereas AAV2-ESGHGYF was detected only in few cells in this compartment (Figure 1f).

Lower but detectable transduction was also observed in several peripheral organs. In the liver, Bovine AAV mediated higher transduction than AAV4 and AAV2-ESGHGYF, both of which showed minimal activity in this tissue (Figure 1c-e). However, liver transduction by Bovine AAV remained substantially lower than that of AAV9, which produced widespread hepatocyte transduction. Bovine AAV-derived GFP signal in the liver was detected mainly in hepatocytes and, less frequently, in endothelial cells (Figure 1c-e).

In the brain, all three capsids primarily targeted capillary and large vessel endothelial cells (Figure 1e), with AAV2-ESGHGYF yielding the highest vector genome and RNA levels (Figure 1c-d).

In the kidney, Bovine AAV and AAV4 both transduced glomerular podocytes and endothelial cells (Figure1e), with Bovine AAV achieving significantly higher DNA and RNA expression than AAV4 (Figure1c-d). AAV9 also transduced the kidney, but with a distinct pattern characterized by prominent expression in glomerular podocytes and renal tubules (Figure 1e). By contrast, AAV2-ESGHGYF showed only minimal renal transduction, limited to a small number of capillaries (Figure 1d-e).

Distinct transduction profiles were also observed in the heart. AAV9 produced the highest RNA and protein expression and predominantly transduced cardiomyocytes (Figure 1d-e). AAV4 also mediated appreciable cardiac transduction, mainly in cardiomyocytes and, to a lesser extent, in endothelial cells (Figure 1d-e). In contrast, AAV2-ESGHGYF transduced vascular endothelial cells selectively, whereas Bovine AAV primarily targeted the endocardium and only occasionally vascular endothelial cells (Figure 1e).

Taken together, Bovine AAV, AAV4, and AAV2-ESGHGYF each showed preferential lung-directed transduction after systemic delivery, with some degree of peripheral organ activity being detected for each vector. Among the three capsids, Bovine AAV was the most effective transducer of both alveolar type I pneumocytes and lung endothelial cells.

### Bovine AAV transduces primary human lung cells more efficiently than conventional AAV serotypes

Although i*n vitro* transduction assays do not fully recapitulate the complexity of *in vivo* organ environments, where factors such as the extracellular matrix, immune responses, vascular delivery, and anatomical structure play crucial roles, they remain useful for assessing vector performance in relevant human primary cell types. Given the robust and selective transduction of mouse pulmonary alveolar epithelial cells and lung microvascular endothelial cells by Bovine AAV *in vivo*, we next evaluated whether Bovine AAV could transduce corresponding primary human lung cell populations *in vitro*.

Primary human pulmonary alveolar epithelial cells, lung microvascular endothelial cells, bronchial tracheal epithelial cells, and small airway epithelial cells were exposed to Bovine AAV, AAV4, AAV2, AAV2-ESGHGYF, or AAV9. Transduction was first assessed 3 days after vector exposure by fluorescence microscopy at 1.0 × 10^6^vg/cell (Figure 2a), and then quantified by flow cytometry across four doses: 1.0 × 10^6^, 5.0 × 10^5^, 2.5 × 10^5^, and 1.25 × 10^5^vg/cell (Figure 2b-c, Supplemental Figure 2a-b).

**Figure 2.**
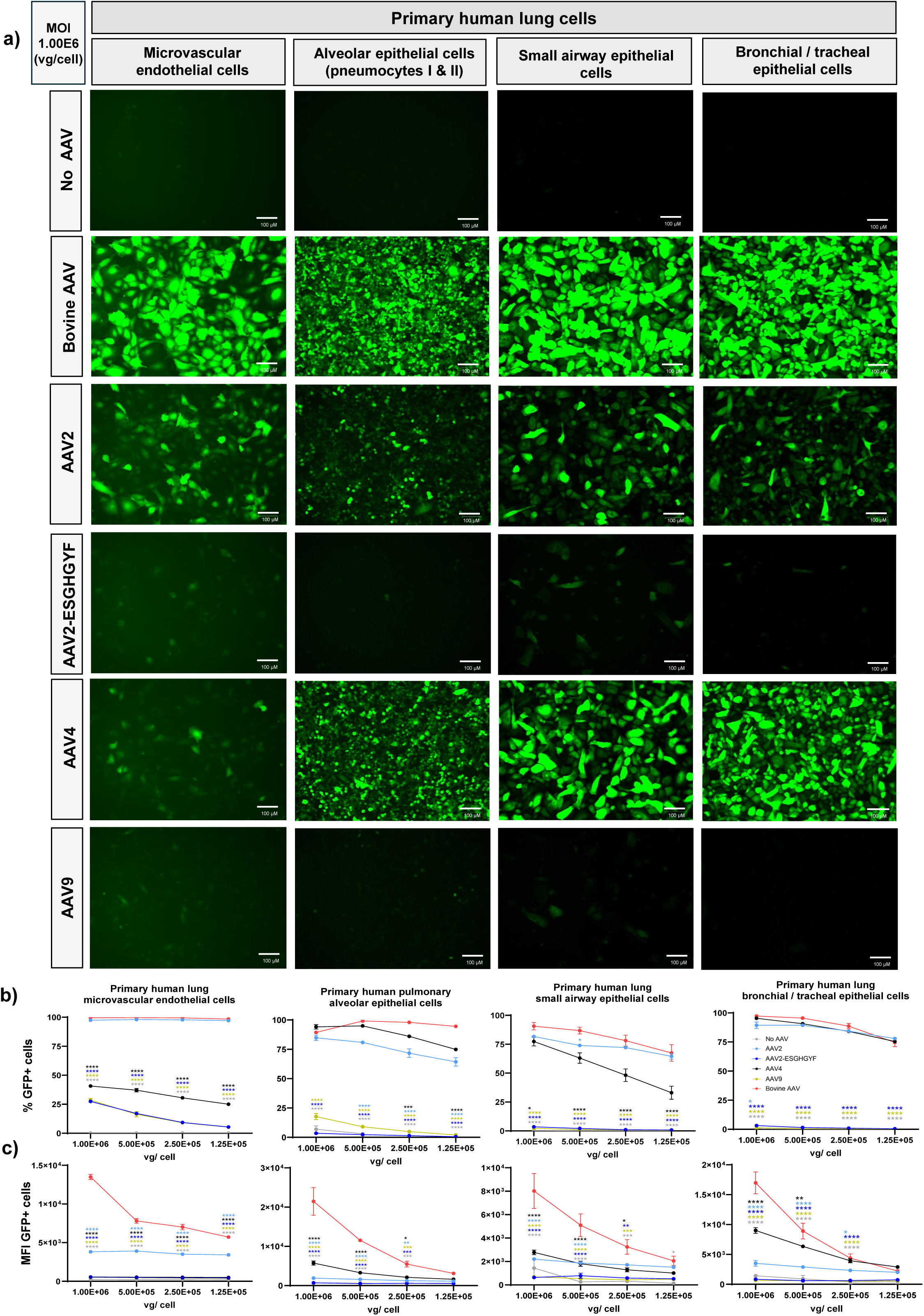
***In vitro* transduction of primary human lung cells with AAV2, AAV2-ESGHGYF, AAV4, AAV9 and Bovine AAV.** Primary human lung cells, including microvascular endothelial cells, alveolar epithelial cells type I and II, small airway epithelial cells, and bronchial tracheal epithelial cells were transduced 24 h after seeding with AAV2, AAV2-ESGHGYF, AAV4, AAV9, or Bovine AAV vectors carrying an ss-CAG-GFP cassette. Vectors were added in duplicates or triplicates (depending on cell availability) at multiplicities of infection (MOI) of 1.0 × 10^6^,5.0 × 10^5^,2.5 × 10^5^,and 1.25 × 10^5^ vg per cell. GFP expression was assessed 3 days after transduction by fluorescence microscopy at 1.0 × 10^6^vg per cell and by flow cytometry at all tested MOIs. A negative control (No AAV) was added and illustrates untransduced cells. *n* = 2 technical replicates for microvascular endothelial cells and alveolar epithelial cells; *n* = 3 technical replicates for small airway epithelial cells and bronchial / tracheal epithelial cells. **a) Representative fluorescence microscopy images** of primary human lung cells 3 days after transduction at 1.0 × 10^6^ vg per cell. Scale bars, 100 μm. **b) Percentage of GFP positive cells** and **c) Median fluorescence intensity of GFP positive cells** in primary human lung cell types quantified by flow cytometry 3 days after transduction across all tested MOIs. Data are shown as mean ± SD of 2-3 technical replicates. Statistical analysis was performed using two-way ANOVA with Dunnett multiple-comparisons test, with each AAV compared to Bovine AAV. Significance is shown relative to Bovine AAV (∗p<0.05, ∗∗p<0.01, ∗∗∗p<0.001, ∗∗∗∗p<0.0001).

Bovine AAV showed robust transduction across all tested primary human lung cell types, including pulmonary alveolar epithelial cells, bronchial tracheal epithelial cells, small airway epithelial cells, and lung microvascular endothelial cells (Figure 2a-c, Supplemental Figure 2b). Across the two highest doses, Bovine AAV consistently produced the highest median fluorescence intensity, and significantly outperformed the other capsids tested (*p* < 0.0001) (Figure 2c). Among the comparator vectors, AAV4 showed comparable transduction to Bovine AAV in pulmonary alveolar epithelial cells, bronchial tracheal and small airway epithelial cells and but remained ineffective in human lung microvascular endothelial cells. AAV2 showed detectable transduction in all cell types (Figure 2a-b), although generally at lower MFI levels than Bovine AAV (Figure 2c). In contrast, AAV2-ESGHGYF and AAV9 showed minimal or undetectable transduction across the primary human lung cell types tested (Figure 2a-c, Supplemental Figure 2b).

To determine whether the liver signal observed for Bovine AAV in mice extended to human hepatocytes, primary human hepatocytes were included in the *in vitro* panel (Supplemental Figure 2c-f). Under these conditions, Bovine AAV did not measurably transduce primary human hepatocytes. Similarly low activity was observed for the other AAV capsids tested, whereas hepatocyte permissiveness was confirmed using AAV6 as a positive control (Supplemental Figure 2c-f).

Overall, Bovine AAV showed the broadest and most consistently strong transduction across the panel of primary human lung-resident cells tested. The strong activity of Bovine AAV in primary human alveolar epithelial cells and lung microvascular endothelial cells highlights its relevance to human pulmonary target cells.

### Bovine AAV demonstrates scalable production and acceptable capsid quality

While vector tropism is a critical parameter for AAV candidate selection, manufacturability is equally essential for technical development. We therefore evaluated the production yield, purification performance, and capsid quality of Bovine AAV at larger scale (2.8L) and compared its performance with that of AAV9, a serotype with well-established manufacturing characteristics ^19,23,24^ (Figure 3, Table S2).

**Figure 3.**
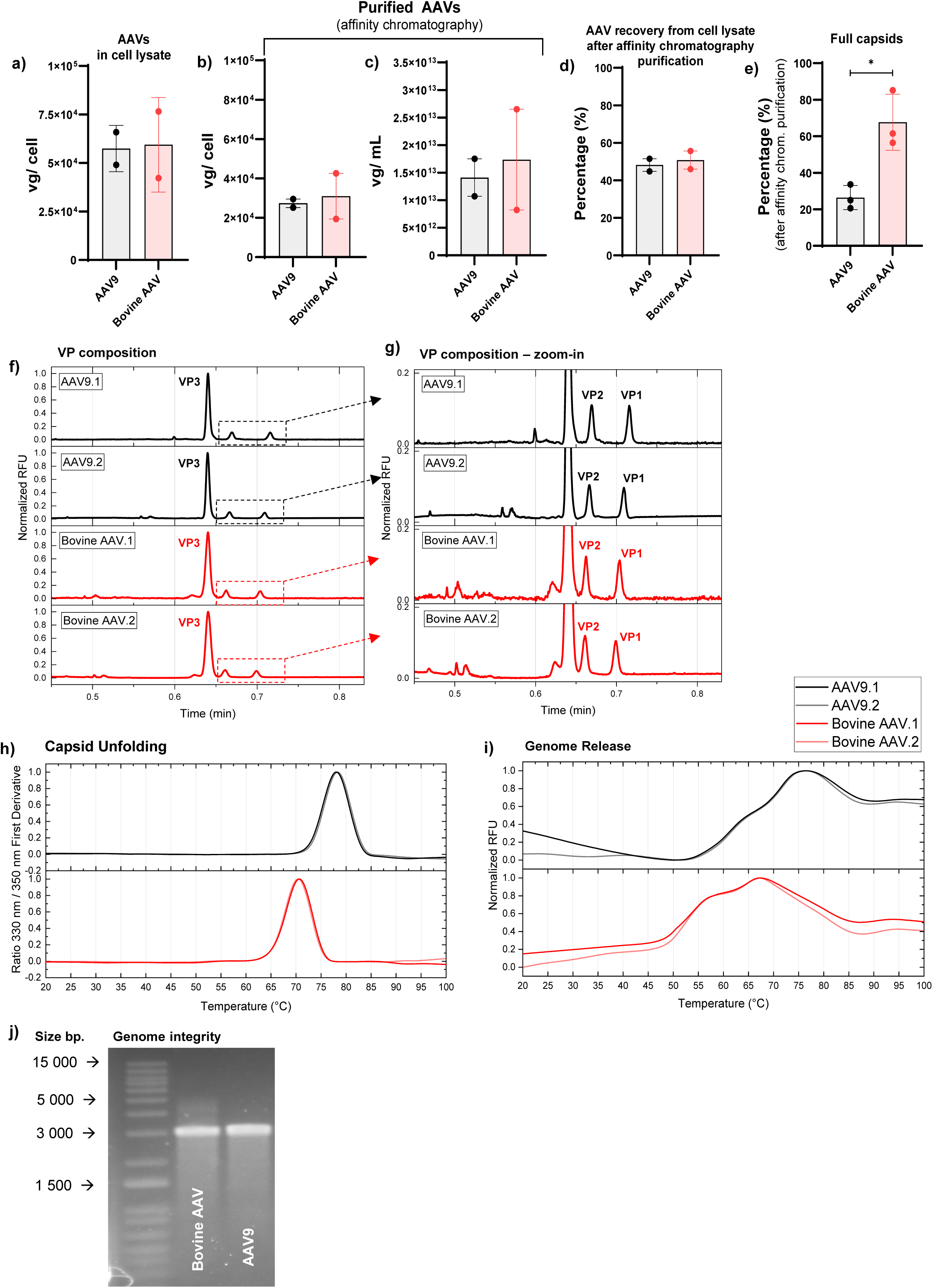
Large-scale production, purification, and biophysical characterization of AAV9 and Bovine AAV. Recombinant AAV9 and Bovine AAV vectors carrying an ss-CAG-GFP genome were produced at large scale in (2.8 liters for Figure 3a-d or 2.8L and 1L for Figure 3e) HEK Expi293 suspension cultures using a three-plasmid transfection system. Viral particles were harvested from cell lysates and purified by affinity chromatography. Full capsids were further enriched by iodixanol gradient ultracentrifugation. Vector genome yields at different production and purification stages were quantified by ddPCR. Biophysical analyses in panels f-i were performed on iodixanol-enriched full capsid preparations (2.8L productions). For a-d), owing to the limited number of independent batches, comparisons are descriptive, therefore data in panels a-d are shown as mean (SD) from two independent production batches per capsid. **a) Vector genome yield in crude cell lysates**, expressed as vector genomes per cell (*vg*/*cell*). Mean (SD) from two independent batches (2.8L productions): AAV9 = 5.75 × 10^4^ (1.20 × 10^4^); Bovine AAV = 5.94 × 10^4^ (2.43 × 10^4^) vg/cell. **b) Vector genome yield in the affinity chromatography-purified fraction**, expressed as vector genomes per cell. Mean (*SD*) from 2 independent batches (2.8L productions): AAV9 = 2.74 × 10^4^(3.04 × 10^3^); Bovine AAV = 3.10 × 10^4^(1.64 × 10^4^) vg/cell. **c) Vector genome titer in the affinity chromatography-purified fraction**, expressed as vector genomes per mL. Mean (*SD*) from 2 independent batches (2.8L productions): AAV9 = 1.41 × 10^13^(4.81 × 10^12^);Bovine AAV = 1.74 × 10^13^(1.29 × 10^13^)vg/mL. **d) Recovery after affinity chromatography purification**, expressed as the percentage of vector genomes recovered from crude cell lysate. Mean (*SD*) from 2 independent batches (2.8L productions): AAV9 = 48.20 (4.76); Bovine AAV = 50.82 (6.86)%. **e) Percentage of full (genome-containing) capsids obtained in the affinity chromatography-purified fraction**. The percentage of full capsids was estimated from the ratio of vector genomes (*vg*/*mL*) to total viral particles (*VP*/*mL*) calculated as (*vg*/*mL* ÷ *VP*/*mL*) × 100. Unpaired t-test (p=0.0128), mean (*SD*) from 3 independent batches (2.8L & 1L productions): AAV9 = 26.39 (6.62); Bovine AAV = 67.68 (15.34); **f) Capsid protein composition by high-throughput capillary gel electrophoresis.** Viral protein composition was analyzed by high-throughput microfluidic chip-based capillary gel electrophoresis under denaturing conditions (HT CE-SDS) resolving VP1, VP2, and VP3. Results from two independent 2.8L production batches are displayed. **g) Expanded view of capsid protein composition.** Zoomed electropherograms highlighting the VP1 and VP2 peaks in AAV9 and Bovine AAV preparations. **h) Capsid unfolding analysis by nano differential scanning fluorimetry.** Thermal unfolding profiles of AAV9 and Bovine AAV capsids are shown as the first derivative of the fluorescence ratio at 330 nm and 350 nm as a function of temperature. Results from two independent 2.8L production batches are displayed for each capsid. **i) Genome release analysis by dye-based differential scanning fluorimetry.** Thermal release of encapsidated genomes was assessed using a SYBR Gold-based DSF assay. Curves are shown as normalized relative fluorescence units as a function of temperature. Results from two independent 2.8L production batches are displayed for each capsid. **j) Genome integrity analysis.** Packaged vector genomes were extracted and analyzed by alkaline agarose gel electrophoresis under denaturing conditions to assess genome integrity. Both AAV9 and Bovine AAV contained a single-stranded GFP genome of 2,983 bases.

**Figure 4.**
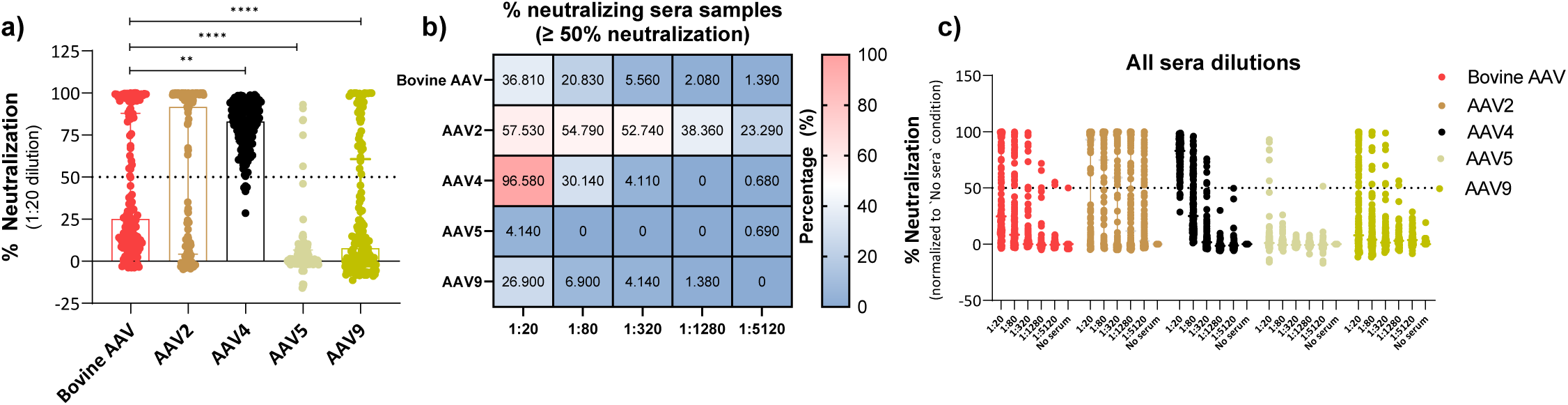
Pre-existing neutralizing antibodies against Bovine AAV, AAV2, AAV4, AAV5, and AAV9 in healthy adult cohorts from the USA and Switzerland. Pre-existing neutralizing antibody responses against Bovine AAV, AAV2, AAV4, AAV5, and AAV9 were assessed in serum samples from healthy adults from the USA (*n* = 73) and Bern, Switzerland (*n* = 72) using a cell-based transduction inhibition assay. Briefly, recombinant AAV vectors carrying an ss-CAG-GFP cassette were incubated with serially diluted human sera (1:20; 1:80; 1:320; 1:1280; 1:5120) for 1 h before addition to HEK293T cells. Transduction efficiency was quantified 3 days later by flow cytometry. Normality was assessed using the Shapiro Wilk test. As the data were not normally distributed, differences in neutralizing antibody seroprevalence across AAV capsids were analyzed using the nonparametric Friedman test followed by Dunn multiple-comparisons test. Statistical significance: ∗*p*<0.05, ∗∗*p*<0.01, ∗∗∗*p*<0.001, ∗∗∗∗*p*<0.0001. **a) Neutralization at the lowest sera dilution.** Percent neutralization at a serum dilution of 1:20, normalized to positive control wells (100% transduction) and negative control wells (0% transduction). Samples were classified as neutralizing if they inhibited transduction by at least 50 percent; the horizontal dashed line indicates this threshold. Each point represents one serum sample, shown as the mean of two technical replicates. Data are presented as median with interquartile range (IQR), 25th to 75th percentile: Bovine AAV (median: 25.19, IQR: 9.91 *to* 87.91; min-max:-3.84 *to* 100.0); AAV4 (median: 83.10, IQR: 71.02 *to* 92.01; min-max: 28.60 *to* 98.92); AAV2 (median: 91.84, IQR: 4.21 *to* 99.30; min-max:-4.81 *to* 99.98); AAV5 (median: 1.09, IQR:-0.81 *to* 6.64; min-max:-16.00 *to* 93.05); AAV9 (median: 7.87, IQR:-3.07 *to* 60.73; min-max:-11.36 *to* 99.98). Statistical differences assessed by Friedman test followed by Dunn multiple-comparisons test: Bovine AAV *vs* AAV4: *p* = 0.036; Bovine *vs* AAV2: *p =* 0.633; Bovine AAV *vs* AAV5 / AAV9: *p*<0.0001. Shown are only statistical differences relative to Bovine AAV. **b) Seroprevalence of neutralizing antibodies across sera dilutions.** Heatmap showing the percentage of sera classified as neutralizing (≥ 50%inhibition of transduction) for each AAV serotype across the five serum dilutions tested in the USA and Bern cohorts. **c) Neutralization across all sera dilutions tested.** Percent neutralization of the indicated AAV capsids measured across the five serum dilutions for all sera samples tested (USA + Switzerland cohorts → 145 sera). Each point represents one serum sample at a given dilution. The horizontal dashed line indicates the 50 percent neutralization threshold, with values above this line considered positive for neutralizing activity.

Recombinant Bovine AAV and AAV9 were produced in 2.8 L HEK Expi293F suspension cultures using a standard three-plasmid transfection system comprising an adenoviral helper plasmid, a packaging plasmid encoding AAV2 Rep and the corresponding Bovine AAV or AAV9 Cap genes, and an ss-CAG-GFP transfer cassette. Viral particles were harvested from cell lysates, purified first by affinity chromatography, and then enriched for genome-containing particles by iodixanol gradient ultracentrifugation.

At 2.8 L scale, Bovine AAV and AAV9 were produced with comparable efficiency, yielding similar vector genome output per cell in crude lysate (Figure 3a). Comparable yields were also observed after affinity purification when measured as both vg/cell and vg/mL (Figure 3b-c). Approximately 50% of vector genomes were recovered after affinity chromatography for both Bovine AAV and AAV9, indicating similar purification performance under the conditions tested (Figure 3d). Analysis of affinity-purified fractions across several independent batches (1L and 2.8L productions) further revealed that Bovine AAV contained a significantly higher proportion of full capsids than AAV9 (Figure 3e).

To further characterize capsid properties relevant to manufacturability and *in vivo* performance, we analyzed capsid quality and stability after ultracentrifugation-based enrichment of genome-containing particles, as this represents the final vector preparation used in downstream pre-clinical and clinical application. Charge detection mass spectrometry (CD-MS) confirmed a higher proportion of full capsids in the analyzed Bovine AAV fraction compared with the corresponding AAV9 fraction (Table S2). Full capsid content varied across collected fractions for both capsids across the two independent experiments, indicating that this attribute is influenced by the UC-related enrichment and fractionation process, rather than being an intrinsic capsid-specific property (Table S2, Supplemental Figure 3a-b). Analysis of capsid protein composition by high-throughput capillary electrophoresis sodium dodecyl sulfate (HT CE-SDS) revealed that Bovine AAV exhibited VP3:VP2:VP1 ratios close to the canonical 10:1:1 stoichiometry, similar to AAV9 (Figure 5f-g; Table S2). Viral genome integrity assessment by alkaline reducing gel electrophoresis demonstrated that both serotypes packaged single-stranded genomes of the expected length (2,983 bp) without evidence of truncation or degradation, confirming intact genome encapsidation (Figure 3j).

**Figure 5.**
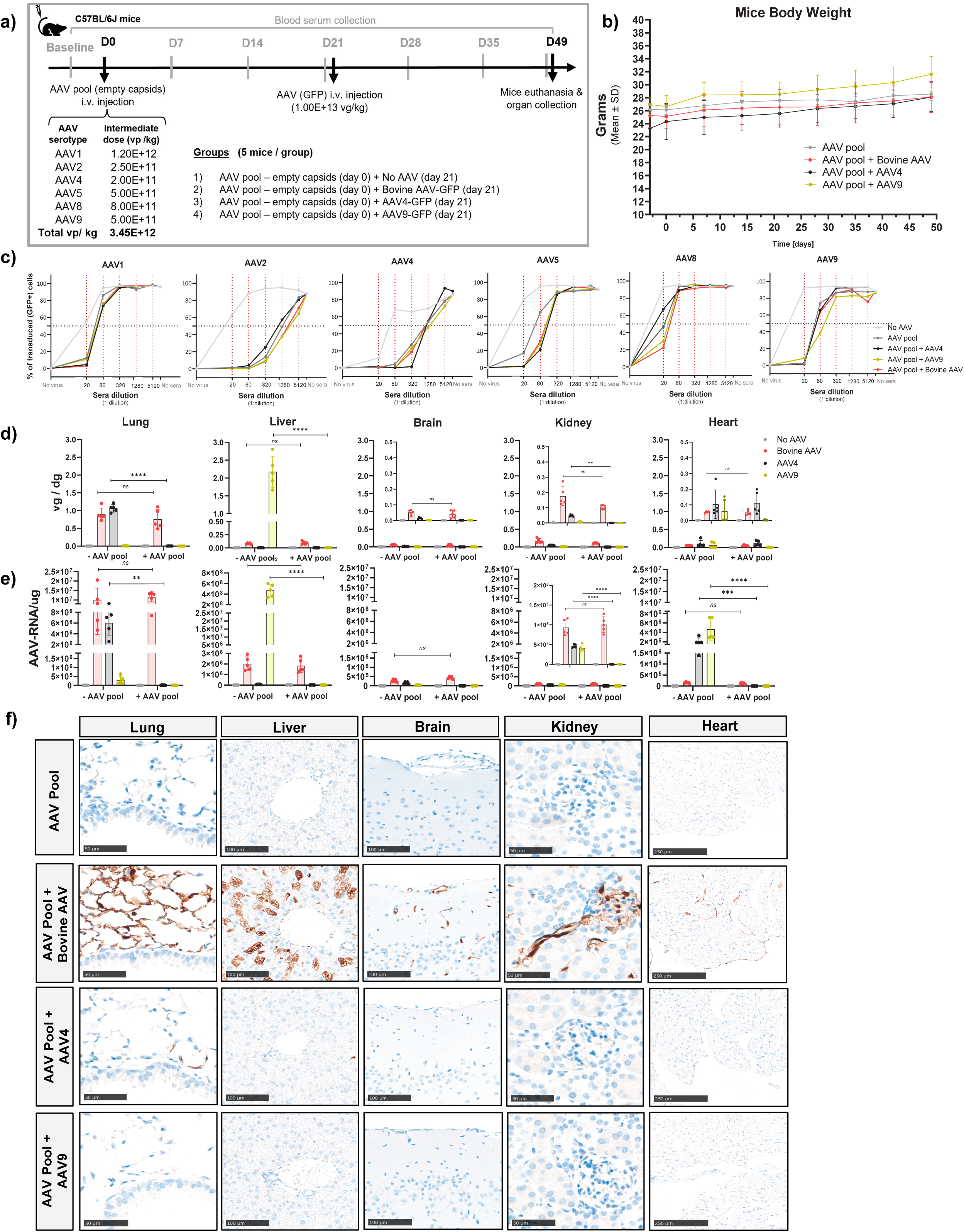
Effect of prior immunization with a pool of human and NHP derived AAV capsids on the *in vivo* transduction efficiency of Bovine AAV, AAV4, and AAV9. **a) Study design.** Male C57BL/6J mice aged 9 to 11 weeks were injected intravenously with a pooled mixture of human and NHP-derived AAV empty capsids comprising AAV1, AAV2, AAV4, AAV5, AAV8, and AAV9 at the indicated doses. Twenty-one days later, mice were assigned to experimental groups (*n* = 5 per group) and injected intravenously with Bovine AAV, AAV4, or AAV9 vectors carrying an ss-CAG-GFP cassette at a dose of 1 × 10^13^vg/kg, corresponding to 2.5 × 10^11^vg per 25 g mouse. One group received AAV formulation buffer at the second injection and served as the pool only control. Four weeks after the second injection, mice were euthanized and lung, liver, brain, kidney, and heart tissues were collected for downstream analyses. Blood was collected from the lateral saphenous vein before AAV pool administration and every 7 days thereafter throughout the 49 day study period for serological analyses. Of note, to minimize the number of animals used, the non-immunized vector-treated comparator groups shown in Figure 1 were generated in parallel using age-and sex-matched mice from the same cohort under identical experimental conditions and are included here, as ‘-AAV pool‘, for direct comparison with the AAV pool-immunized groups (+ AAV pool). **b) Body weight monitoring.** Body weight was recorded 3 days before AAV pool administration and every 7 days throughout the study. Data are shown as mean (SD) body weight in grams for *n* = 5 mice per group. Differences in body weight changes over time depending on the injected AAVs were analyzed using a two-way repeated-measures ANOVA with AAV treatment group as the between-subject factor and time as the within-subject factor, followed by Sidak multiple-comparisons test. **c) Evaluation of humoral immunity induced by AAV pool administration - Neutralizing activity against the AAV capsids included in the pool.** Neutralizing activity against AAV1, AAV2, AAV4, AAV5, AAV8, and AAV9 was assessed using a cell based transduction inhibition assay. For each experimental group, and each mouse respectively, sera collected at days 14 and 21 post immunization were pooled and serially diluted (1:20; 1:80; 1:320; 1:1,280; 1:5,120) before incubation with the corresponding GFP expressing AAV reporter vector for 1 h at 37 °C. The serum vector mixtures were then added to HEK-293T cells, and transduction efficiency was quantified 72 h later by flow cytometry as the percentage of GFP positive cells. Vectors were used at MOIs previously determined to yield approximately 80 to 90 percent transduction in the absence of serum (Supplemental Figure 5e). Wells containing cells plus virus without serum and cells without virus or serum were included as positive and negative controls, respectively, for normalization of transduction efficiency. Sera were classified as neutralizing if they reduced transduction by at least 50 percent at the lowest dilution tested (1:20). The dashed horizontal line indicates the 50 percent neutralization threshold. Points represent the mean measured transduction value for each group at the indicated serum dilutions. **d) Tissue vector biodistribution** and e**) Transgene expression.** Recombinant AAV vector genomes were quantified by ddPCR and are presented as vector genomes per diploid genome (*vg* /*dg*), whereas transgene expression was quantified by RT-ddPCR and is presented as AAV RNA per microgram of tissue. Because expression levels in the brain, kidney, and heart were lower than in the other organs, enlarged views of these tissues are included to facilitate comparison among groups. Points represent individual mice and bars indicate mean (SD). Statistical significance was assessed by two-way ANOVA with Sidak multiple comparisons test (∗p<0.05, ∗∗p<0.01, ∗∗∗p<0.001, ∗∗∗∗p<0.0001). **h) Representative histology images.** Representative immunohistochemistry images of FFPE sections from lung, liver, brain, kidney, and heart illustrate localized transgene expression following administration of Bovine AAV, AAV4, or AAV9 in mice previously immunized with the AAV capsid pool. Scale bars, 50 μm for lung and kidney, 100 μm for liver and brain, and 250 μm for heart.

Thermal stability analysis revealed that Bovine AAV had capsid unfolding temperatures which were 7-10°C lower than AAV9, indicating slightly reduced structural robustness of the capsid shell (Figure 3h, Table S2). Bovine AAV also exhibited a 9-10°C lower onset temperature for genome release, suggesting that genome extrusion occurs at a lower thermal threshold (Figure 3i, Table S2).

Collectively, these data indicate that Bovine AAV can be produced and purified efficiently at larger scale, with vector yield, quality, and genome integrity comparable to well-established AAV serotypes. The narrower thermal stability margins may have implications for large-scale production, formulation, storage, and transportation, and needs to be considered during future process optimization.

### In healthy individuals, Bovine AAV displays an intermediate seroprevalence profile relative to established AAV serotypes

Because pre-existing humoral immunity can substantially limit eligibility for AAV-mediated gene therapy, we next examined the prevalence of neutralizing antibodies against Bovine AAV in healthy adults and compared it with selected human and NHP-derived AAV serotypes. Bovine AAV was evaluated alongside AAV2, the serotype with the highest seroprevalence worldwide ^25,26^, AAV4, a key comparator for lung transduction, AAV5, noted for the lowest neutralizing antibody seroprevalence in both adult and pediatric populations ^6^, and AAV9, which typically shows intermediate neutralizing antibody prevalence in healthy adults ^6^. To the best of our knowledge, this study represents the first investigation of pre-existing immunity to Bovine AAV in healthy individuals.

Serum samples were obtained from three independent cohorts of healthy adults from the United States of America (USA) and Switzerland. In total, 73 samples were collected from healthy individuals in the USA (mean age 48.9 years, SD 15.5), from two independent commercial sources. This cohort included 42 males (mean age: 40.6 years, SD 14) and 31 females (mean age: 41.2 years, SD 13.9) and encompassed multiple self-reported ethnic backgrounds, including White, Black, Asian, and other racial groups (Figure 4, Table 2). Additional 72 samples were obtained from healthy individuals in Bern, Switzerland, through the Novartis sample donation program (mean age 46.8 years, SD 16.1 where available). This cohort included 36 males (mean age: 49.6 years, SD 15.7) and 20 females (mean age: 46.7 years, SD 15.5). Ethnicity data was not recorded for the Swiss cohort, and sex and age information were unavailable for 16 individuals within this cohort (Table 2). Neutralizing activity against each capsid was measured using *in vitro* transduction inhibition assays across five serial serum dilutions. For each serotype, the multiplicity of infection required to transduce 80-90% of cells was firstly determined (Supplemental Figure 5e and 6b). Sera were considered neutralizing if at least 50% of vector transduction was blocked at the lowest sera dilution tested, namely 1:20. The full distribution of neutralizing responses across all dilutions is shown in Figure 4c and Supplemental Figure 5c, 5f. Inter-serotype comparisons were performed using non-parametric matched analyses as described in the methods section (Statistics; Supplemental Figure 5g-h).

Bovine AAV exhibited an intermediate neutralization profile relative to the comparator capsids. At the lowest serum dilution tested, 1:20, 37% of samples neutralized Bovine AAV (Figure 4a-c). Across the dilution series, Bovine AAV was neutralized less frequently than AAV2 and AAV4, including significantly lower neutralization than AAV4 at both 1:20 and 1:80, but more frequently than AAV5 and AAV9 (Figure 4a-c). Among the serotypes tested, AAV4 showed the highest frequency of neutralization at the lowest serum dilution, with 100% of samples scoring positive at 1:20 (Figure 4a-b). However, this response declined sharply at higher dilutions, dropping to 30% at 1:80 and 4% at 1:320, indicating the presence of low-titer neutralizing activity that is detectable only at low serum dilution (Figure 4a-b). In contrast, AAV2 showed less frequent but more persistent neutralization across the dilution series, with 57.5% of samples neutralizing at 1:20 and 23.3% remaining positive at 1:5120 (Figure 4b). AAV5 showed the lowest neutralization rates across all dilutions, with 5.4% neutralizing at 1:20 and no detectable neutralization at higher dilutions (Figure 4b). AAV9 displayed low to intermediate neutralization across the cohort (Figure 4a-c).

Because AAV seroprevalence can vary across geographic regions (Chhabra et al., 2024), the two cohorts were also analyzed separately (Supplemental Figure 4). Similar serological patterns were observed in both the USA (Supplemental Figure 4a-c) and the Swiss cohorts (Supplemental Figure 4d-f), with Bovine AAV retaining an intermediate profile relative to comparator capsids. Notably, the Swiss cohort showed a lower number of neutralizing sera against Bovine AAV, AAV2 and AAV9 than the USA cohort (Supplemental Figure 4e-f).

Overall, these serological analyses identify Bovine AAV as a capsid with intermediate pre-existing humoral immunity in healthy adults, lower than AAV2 and AAV4 but higher than AAV5 and AAV9.

### Bovine AAV efficiently transduces the lungs *in vivo* despite pre-existing immunity to multiple AAV serotypes

There is often considerable sequence and structural similarities between different AAVs ^27^, therefore antibodies raised against one serotype may cross-react with related capsids, thereby reducing the effectiveness of systemic gene therapy and complicating gene re-administration with alternative capsids ^28–30^. Despite that Bovine AAV is phylogenetically distinct from commonly used human and NHP-derived AAVs, we investigated whether prior exposure to a panel of these capsids would affect Bovine AAV-mediated transduction *in vivo*.

To establish a model of pre-existing anti-AAV immunity, male C57BL/6J mice were immunized intravenously with a pool of empty capsids comprising AAV1, AAV2, AAV4, AAV5, AAV8, and AAV9 and blood samples were collected weekly for serological analysis (Supplemental Figure 5a). Preliminary dose-finding experiments were performed to identify doses that elicit moderate humoral responses for each serotype, defined by intermediate anti-AAV IgG/ IgM and neutralizing antibody levels, to mimic a pre-existing exposure to naturally occurring AAVs in humans. Mice receiving the pooled capsids were monitored throughout the 49-day study and no significant changes in body weight were observed relative to non-immunized controls (Supplemental Figure 5b).

Serological analysis showed that anti-AAV IgM responses increased transiently after immunization, peaking early and declining thereafter (Supplemental Figure 5c). Background signal was observed in both control and uncoated wells (Supplemental Figure 5c), likely reflecting the polyreactive, non-specific binding properties of pentameric IgM, previously observed in the literature ^31,32^. For most serotypes, the intermediate dose was sufficient to induce stable anti-AAV IgG responses, detectable by days 14–21 and maintained throughout the study period; only AAV1 and AAV4 required dose adjustment (Supplemental Figure 5d). Thus, the optimized final pool formulation included the lower dose of AAV4 and a higher dose of AAV1 (Figure 5a).

To assess the neutralizing capacity of the induced antibodies, we made use of the *in vitro* transduction inhibition assay based on HEK-293T cells (Supplemental Figure 5f). For each serotype, the multiplicity of infection required to transduce 80-90% of cells was firstly determined (Supplemental Figure 5e). Sera collected on days 14 and 21 were pooled together (because of volume limitations) and tested for inhibition of transduction (Supplemental Figure 5f). All serotypes in the pooled intermediate-dose formulation induced neutralizing activity at the lowest serum dilution (Supplemental Figure 5f).

We next evaluated whether this pre-established immunity affected Bovine AAV-mediated transduction *in vivo* (Figure 5a-f). Male C57BL/6J mice were firstly immunized intravenously with the pooled empty capsids at a total dose of 3.45 × 10^12^vg/kg (Figure 5a). Three weeks later, mice received a second intravenous injection of Bovine AAV-GFP at 1.0 × 10^13^vg/kg. Control groups received the same immunization followed by AAV4-GFP, AAV9-GFP, or no second vector (Figure 5a). Four weeks after the second injection, lung, liver, brain, kidney, and heart were harvested for analysis of vector genome abundance, transgene RNA expression, and GFP protein expression (Figure 5d-f). To enable direct comparison of transduction outcomes in the presence and absence of prior AAV pool immunization while minimizing the number of animal used, data from the non-immunized vector-treated groups (- AAV pool) shown in Figure 1 were analyzed alongside the immunized groups (+AAV pool) in Figure 5. These experiments were performed in parallel using age-and sex-matched mice from the same cohort under identical experimental conditions.

All treatment regimens were well tolerated, and no significant body weight changes were observed during either the immunization or challenge phases (Figure 5b). Baseline anti-AAV IgG, IgM, and neutralizing antibody levels were low before vector administration (Figure 5c, Supplemental Figure 6a-b). Following pool immunization, all groups developed intermediate to high anti-AAV IgG and IgM titers against the six capsids included in the formulation (Supplemental Figure 6a-b). As expected, administration of AAV4, AAV9, or Bovine AAV as the second vector further increased corresponding IgG antibody responses (Supplemental Figure 6b). Neutralizing activity against all serotypes present in the pool was detected in sera collected at days 14 and 21 at a 1:20 dilution (Figure 5c).

Having confirmed neutralizing activity against all serotypes included in the pool, we next tested whether sera from AAV pool-immunized mice could cross-neutralize Bovine AAV using the same cell-based transduction inhibition assay (Supplemental Figure 5e, Supplemental Figure 6c). Sera from buffer-injected mice served as negative controls, whereas sera from mice immunized with the AAV pool and subsequently exposed to Bovine AAV served as positive controls. No Bovine AAV neutralization was detected with sera from AAV pool-immunized mice, indicating that antibodies raised against the pooled human and non-human primate-derived AAV capsids did not measurably cross-neutralize Bovine AAV (Supplemental Figure 6c).

Despite the presence of robust humoral immunity against multiple human and non-human primate-derived AAV capsids and consistent with the absence of detectable *in vitro* cross-neutralization, Bovine AAV retained efficient tissue transduction in AAV pool-immunized mice (Figure 5d-f). No significant differences were observed between the Bovine AAV-only group and the AAV pool + Bovine AAV group in vector genome abundance, transgene RNA expression, or GFP protein expression in any tissue analyzed (Figure 5d-f). In contrast, transduction was completely blocked in animals that received the AAV pool followed by a second injection of either AAV4 or AAV9, demonstrating that even low-dose priming is sufficient to prevent re-administration with the same capsid (Figure 5d-f).

Because Bovine AAV shares the greatest capsid sequence similarity with AAV4 (Supplemental Figure 1, Table S1, and Schmidt et al., 2004), we further examined whether AAV4-specific antibodies could cross-neutralize Bovine AAV (Supplemental Figure 6d). Sera from mice immunized with AAV4 alone at doses higher than that included in the pool (5.0 × 10^11^and 5.0 × 10^12^vg/kg) showed no detectable Bovine AAV neutralization (Supplemental Figure 6d), consistent with previous reports (Schmidt et al., 2004).

Together, these findings show that Bovine AAV-mediated transduction is preserved in the presence of pre-existing immunity to a panel of commonly used human and NHP-derived AAV serotypes and is not detectably cross-neutralized by antibodies elicited against these capsids.

## Discussion

Despite encouraging preclinical findings, AAV-based gene therapy for inherited lung diseases has not yet achieved meaningful clinical efficacy. The large surface area of the lung, its multiple physical and biological barriers, pre-existing immunity to human and NHP-derived AAVs, suboptimal transduction efficiency, and rapid cellular turnover have all hindered effective gene delivery by recombinant AAV vectors ^3^. Several natural serotypes, including AAV1, AAV2, AAV5, AAV6, and AAV9, have been explored preclinically, and AAV1, AAV2, and AAVrh.10 have advanced to clinical testing ^1^. However, these approaches have thus far been limited by insufficient therapeutic efficacy and, in some cases, by pre-existing immunity in the human population.

One strategy to overcome these barriers is to evaluate more evolutionary divergent capsids. Because phylogenetically distant AAVs often retain the ability to enter mammalian cells through distinct receptor interactions, such serotypes may display unique transduction properties in human tissues. Bovine AAV is phylogenetically distant from most human and non-human primate-derived (NHP) AAVs and has previously been reported to exhibit lung-tropic properties ^15,16^. Bovine AAV shares 76% capsid sequence homology with AAV4 ^14^, one of the most structurally distinct AAV serotypes ^17^ and a key comparator in the context of pulmonary delivery. Recent work identified AAV4 as one of the strongest naturally occurring lung-tropic capsids described to date in both mice and NHPs, while also showing relative liver detargeting ^18^. In this context, Bovine AAV becomes particularly interesting to investigate both as a lung-directed vector and as a potentially antigenically distinct alternative to established human and NHP-derived capsids.

In this study, we comprehensively evaluated Bovine AAV as a candidate immune-orthogonal vector for systemic lung-directed gene therapy. To define its strengths and limitations relative to established human and NHP-derived AAVs, including AAV4, we assessed Bovine AAV *in vivo* transduction profile, cross-reactivity with other serotypes, ability to transduce relevant primary human lung cells, seroprevalence in healthy adults, and manufacturability. Together, these parameters provide a framework for evaluating both the biological performance and translational feasibility of Bovine AAV. Intravenous administration of Bovine AAV resulted in robust lung transduction in mice, reaching levels comparable to AAV4 and markedly exceeding those of the engineered lung-tropic vector AAV2-ESGHGYF. Both Bovine AAV and AAV4 showed preferential lung tropism, with the highest transgene expression in the lung, and both maintained detectable transgene expression four weeks after injection. These findings identify Bovine AAV as a strong lung-tropic vector following systemic delivery and place it among the most effective naturally occurring capsids currently described for pulmonary gene transfer in mice.

Our data help contextualize prior studies of Bovine AAV in the respiratory tract. Earlier work using the same Bovine AAV construct reported that intranasal transduction required inhibition of viral transcytosis with tannic acid and was transient, becoming undetectable by eight weeks after administration ^15^. In contrast, a more recent study using the closely related AAV.Bov.Guelph capsid, which differs by only two amino acids, demonstrated robust and durable lung transduction after intranasal delivery without transcytosis inhibition ^16^. Together with the present intravenous findings, these observations suggest that Bovine AAV-like capsids have a broader pulmonary delivery potential than initially appreciated, but that their performance strongly depends on capsid sequence, route of administration and the biological barriers encountered at the airway versus vascular interface.

Vector delivery route is known to influence initial capsid-cell interactions, receptor accessibility, clearance dynamics and transduction distribution ^33–35^. While intranasal delivery must overcome mucociliary clearance and epithelial barriers ^3,36^, often resulting in a heterogeneous airway facing distribution ^20^, intravenous administration provides vascular access to more distal lung compartments and may support a broader distribution across the lung ^37^. Consistent with this, both Bovine AAV and AAV4 primarily transduced pulmonary microvascular endothelial cells and alveolar-region cells *in vivo*, including type I pneumocytes, with little evidence of airway epithelial transduction. Importantly, neither vector detectably transduced type II pneumocytes. Complementary studies in primary human cells further refined this comparison: while both Bovine AAV and AAV4 transduced primary human alveolar epithelial cells, only Bovine AAV efficiently transduced primary human lung microvascular endothelial cells. This distinction may be especially relevant translationally, as it suggests that Bovine AAV may better preserve its endothelial tropism across species and may therefore offer advantages for pulmonary vascular targeting.

This transduction pattern has important implications for therapeutic positioning. In its current form and through systemic delivery, Bovine AAV appears more suited for applications involving the pulmonary vasculature than for diseases requiring broad correction of the airway epithelium or alveolar type II cells. In particular, its preferential transduction of lung endothelial cells supports further evaluation in diseases such as pulmonary arterial hypertension (PAH), a progressive pulmonary vascular condition characterized by endothelial dysfunction, vascular remodeling, increased pulmonary vascular resistance, and ultimately right heart failure ^38^. Notably, AAV-based gene transfer has already shown therapeutic benefit in preclinical models of pulmonary hypertension, including studies using AAV1-SERCA2a that reduced pulmonary vascular remodeling and improved hemodynamic parameters ^39^. Because endothelial injury and dysregulated vascular signaling are central to PAH pathogenesis ^40,41^, vectors with efficient access to the pulmonary endothelium may be especially valuable for gene replacement or pathway-modulating approaches, including those involving BMPR2 or SERCA2a-related signaling ^39,42,43^. Conversely, Bovine AAV appears less suitable, at least in its current form, for disorders that require direct correction of alveolar type II pneumocytes. This limitation argues against immediate application of Bovine AAV for surfactant-related disorders such as SFTPB, SFTPC, and ABCA3 deficiency, for which efficient targeting of type II pneumocytes would likely be essential ^44–46^. For such indications, additional capsid engineering or alternative delivery approaches would be required. While both Bovine AAV and AAV4 showed clear preferential lung expression, neither vector was fully lung-restricted, as expected with systemic administration. For Bovine AAV, significantly lower but detectable expression in the liver, heart, kidney, and brain indicates some off-target distribution and transduction. AAV4 also showed multi-organ activity, with particularly notable cardiac expression, consistent with previous reports ^18,21,47,48^. Because such broad tropism may complicate therapeutic applications, this limitation could be mitigated through the use of lung cell type-specific promoters (e.g., SPC, AGER, CC10, FOXJ1, VE-cadherin) ^49–53^, microRNA-mediated detargeting (e.g., miR-122 for liver^54^, 2010 or miR-1 for muscle/heart ^55,56^ or additional capsid engineering to further restrict off-target expression ^5,37,57^.

Of translational relevance, Bovine AAV retained efficient *in vivo* transduction in mice previously immunized with a pool of human and NHP-derived AAV capsids, including AAV4, despite the development of robust humoral immunity to all serotypes in the pool. Consistent with this observation, sera from AAV pool-immunized mice did not neutralize Bovine AAV *in vitro*, indicating that antibodies elicited by these conventional capsids did not show detectable functional cross-neutralization of Bovine AAV. These findings extend earlier *in vitro* observations showing that Bovine AAV is not neutralized by sera from mice immunized with either AAV2, AAV4, or AAV5 ^14^. This property is relevant for systemic gene therapy, where pre-existing immunity to common human AAV serotypes can exclude a substantial fraction of patients and complicate sequential gene re-administration through a different capsid. The absence of detectable cross-neutralization between Bovine AAV and AAV4 is particularly notable given their close phylogenetic relationship and shared pulmonary tropism, and suggests that, in patients lacking neutralizing antibodies to both capsids, gene re-administration could in principle be achieved by delivering the same transgene first with AAV4 and subsequently with Bovine AAV or *vice versa*.

In addition to efficient transduction of target cells, serological profiling is an essential component of vector evaluation in AAV-based gene therapy, considering that pre-existing humoral immunity is a major determinant of patient eligibility and therapeutic efficacy. Bovine AAV displayed an intermediate neutralization profile in healthy adults, with lower neutralization than AAV2 and AAV4 but higher neutralization than AAV5 and AAV9. This profile is particularly relevant in comparison to AAV4, which showed strong pulmonary transduction but the highest neutralization frequency. By contrast, Bovine AAV achieved similarly strong lung-directed activity *in vivo* while being less frequently neutralized in human sera, suggesting that it may retain utility in a broader subset of individuals. Thus, although Bovine AAV is not immunologically silent, it may offer a more favorable balance between lung tropism and pre-existing humoral immunity than AAV4.

The lower frequency of Bovine AAV-neutralizing antibodies observed in the Swiss cohort further suggests that Bovine AAV seroreactivity may vary geographically, consistent with the regional heterogeneity reported for other AAV serotypes ^6^. Although confirmation in larger and more diverse populations is needed, this finding underscores the importance of evaluating candidate vectors across diverse populations during translational development.

In addition to its biological performance and immune profile, Bovine AAV demonstrated favorable manufacturability at larger scale, with high production yields, efficient genome packaging, and a high proportion of full capsids. This is an important translational advantage, as manufacturability is often a key determinant of whether an AAV vector can progress beyond preclinical evaluation. Many capsids fail not because of insufficient biological activity, but because they cannot be produced robustly at the scale and quality required for clinical use ^8,58^. In this regard, Bovine AAV compares favorably with AAV9, a well-established manufacturing benchmark, and appears suitable for downstream process development. This is particularly relevant for pulmonary gene therapy, where the size and complexity of the lung may require substantial vector quantities to achieve effective target coverage. Nevertheless, Bovine AAV exhibited a narrower thermal stability margin compared to AAV9. While the lower genome-release threshold may help the capsid more readily uncoat intracellularly and transduce cells with more efficiency, from a manufacturing standpoint, it may increase susceptibility to changes in full-capsid content under thermal or process-related stress, highlighting the importance of formulation and process optimization.

### Future directions and conclusions

While our findings establish Bovine AAV as a promising lung-tropic and immune-distinct vector, several questions remain for future work. Because all *in vivo* studies were performed in mice, evaluation in non-human primates might be essential to determine whether the lung selectivity and immune-evasion properties observed here are conserved in larger species with airway and endothelial biology more similar to humans. Longer-term studies will also be needed to assess the durability of expression and to exclude delayed immune effects. In addition, disease-specific contexts, such as inflammation, fibrosis, or vascular remodeling, may alter vector access^62,63^ and should be examined to define therapeutic applicability. Despite these considerations, Bovine AAV combines features rarely aligned within a single platform: robust lung transduction *in vivo*, preserved efficacy in the presence of high titers against conventional AAVs, transduction of multiple primary human lung cell types, and scalable manufacturability. Together, these attributes position Bovine AAV as a promising translational candidate and a valuable scaffold for future capsid engineering.

## Material and Methods

### Statistics

The following statistical tests were used to determine the statistically significant differences between the variable assessed: **Body weight measurements following AAV administration:** Figures 1b, Figure 5b, Supplemental Figure 5b→ two-way repeated-measures analysis of variance (ANOVA) with AAV treatment group as the between-subject factor and time as the within-subject factor, followed by Sidak multiple-comparisons test. **Tissue vector biodistribution and transgene expression**: Figure 1c-d → one-way ANOVA with Tukey‘s multiple comparisons test; Figure 5e-f→ two-way ANOVA with Sidak multiple comparisons test. ***In vitro* transduction of primary human lung cells and primary human hepatocytes**: Figure 2b-c, Supplemental Figure 2e-f → two-way ANOVA with Dunnett multiple-comparisons test. **Percentage of full capsids**: Figure 3e → unpaired t-test. **Pre-existing immunity to Bovine AAV and comparator capsids in the human population:** Figure 4a, Supplemental Figure 5a and 5d → Shapiro-Wilk test and normal Q-Q plots were used to assess the normal distribution of the neutralization data. Because data showed significant deviation from normality (p < 0.010) and skewness and ceiling effects were observed (Supplemental Figure 4g-h), the non-parametric Friedman’s test followed by Dunn’s multiple comparison post hoc tests were used for inter-serotype comparisons. Significance is reported as following: ∗p<0.05, ∗∗p<0.01, ∗∗∗p<0.001, ∗∗∗∗p<0.0001. Unless otherwise stated, all quantitative data are presented as means (standard deviation). All statistical analyses were performed using GraphPad Prism 10 software.

### VP1 amino acid sequence similarity assessment between distant and human / NHP-derived AAVs

A total of 25 VP1 amino acid sequences corresponding to the following parvoviruses (PVs) were analyzed (accession numbers are provided in parenthesis): AAAV.DA-1 (NC_006263), AAV1 (NC_002077), AAV2 (J01901.1), AAV3 (NC_001729), AAV3b (AF028705), AAV4 (NC_001829), AAV5 (NC_006152), AAV6 (AF028704), AAV7 (NC_006260), AAV8 (NC_006261), AAV9 (AY530579), AAV10 (AY631965), AAV11 (AY631966), AAV12 (DQ813647), AAV13 (EU285562), AAVrh.10 (AY243015), AAVrh.74 (KH123010), Bovine AAV (AY388617), Bearded Dragon Parvovirus (BDPV) (KP733794), Bat AAV (BtAAV) (NC_014468), Californian Sea Lion (Csl)-AAV1 (NC_038539), Duck Parvovirus (DPV) (NC_006147), DPV_G018-19 (MW380869), Goose Parvovirus (GPV) (NC_001701), and Snake AAV (SAAV) (NC_006148).

VP1 protein sequences were aligned using the MUSCLE algorithm (https://doi.org/10.1093/nar/gkh340) implemented in Geneious Prime 2025. The resulting multiple sequence alignment was used to construct a phylogenetic tree based on genetic distances calculated with the Jukes-Cantor substitution model (http://dx.doi.org/10.1016/B978-1-4832-3211-9.50009-7). Tree inference was performed using the neighbor-joining method (https://doi.org/10.1093/oxfordjournals.molbev.a040454). To assess the robustness of the inferred topology, bootstrap resampling with 1,000 replicates (https://doi.org/10.1111/j.1558-5646.1985.tb00420.x) was applied. All sequence analyses and visualizations were conducted in Geneious Prime 2025.

### AAV production, purification and full-capsid enrichment

All recombinant AAV vectors used in this study were produced in suspension-adapted Expi293F HEK cells, using a transient three-plasmid transfection system. Cells were expanded in chemically defined HEK293 medium (Irvine Scientific) under standard suspension culture conditions and transfected at high viability with an adenoviral helper plasmid, a Rep-Cap plasmid encoding AAV2 Rep and the appropriate serotype-specific Cap gene (see Table S3), and an ss-CAG-GFP transfer plasmid. Plasmids were mixed at an equimolar ratio and complexed with a commercially available AAV transfection reagent (Polyplus) in reduced medium following an internally optimized protocol. Vectors were harvested 72-hours post-transfection from cell lysates. Residual nucleic acids were digested by nuclease treatment, and cells were lysed under detergent-based conditions optimized for recovery of intracellular AAV particles while preserving capsid integrity. Crude lysates were clarified by centrifugation and filtration, treated to terminate nuclease activity, and either processed immediately or stored under controlled conditions before purification.

Clarified AAV-containing lysates were purified by affinity chromatography. Following column loading and washing, bound vectors were eluted under low-pH conditions and immediately neutralized. AAV-containing fractions were pooled and buffer-exchanged into formulation buffer. For full-particle enrichment, affinity-purified vectors were subjected to iodixanol density gradient ultracentrifugation. Fractions enriched for genome-containing capsids were collected, buffer-exchanged, concentrated, sterile filtered, aliquoted, and stored at −80^∘^C until further use.

### Endotoxin level measurement in AAV samples

All recombinant AAV preparations intended for *in vivo* studies were tested for endotoxin contamination using a Limulus amebocyte lysate (LAL) assay. AAV samples were diluted 1:20 in endotoxin-free water and loaded into Endosafe® LAL cartridges (Charles River Laboratories, Cat. No. PTS20F; sensitivity range 0.1–1.0 EU/mL). Endotoxin levels were measured in quadruplicate using an Endosafe® nexgen-PTS™ system (Charles River Laboratories, Cat. No. PTS150K) according to the manufacturer’s instructions and values were reported as EU/mL of final formulation. Only rAAV preparations with endotoxin levels below 0.2 EU/mL were considered acceptable and used for *in vivo* administration.

### AAV genome titration by digital droplet polymerase chain reaction

Recombinant AAV genome copy numbers (titers) were quantified by droplet digital polymerase chain reaction (ddPCR) following AAV production and purification. AAV samples (10 µL) were initially diluted 1:10 in 90 µL of dilution buffer consisting of nuclease-free water (Invitrogen, Cat. No. AM9937), 1× PCR Buffer II (Applied Biosystems, Cat. No. 4379878), 2.5 mM MgCl₂ (Applied Biosystems, Cat. No. 4379878), 0.05% Pluronic F-68 (Gibco, Cat. No. 24040032), and 0.002 mg/mL sheared salmon sperm DNA (Invitrogen, Cat. No. AM9680).

To eliminate contaminating free DNA, 10 µL of the first dilution were transferred into 90 µL of DNase I digestion mix containing 79 µL nuclease-free water, 10 µL 10× DNase buffer, and 1 µL DNase I (2 U/µL; Invitrogen, Cat. No. AM3333), resulting in a 1:100 dilution. Samples were incubated for 30 min at 37 °C in a Bio-Rad C1000 Touch thermocycler (Bio-Rad Laboratories).

To disrupt AAV capsids and allow access to encapsidated genomes, 50 µL of DNase-treated sample were mixed with 50 µL of proteinase K (PK) reaction mix containing 45 µL 2× PK buffer (composed of nuclease-free water, 1% SDS (Invitrogen, Cat. No. AM9822), 0.1 M EDTA (Invitrogen, Cat. No. 15575020), and 0.1 M Tris-HCl, pH 8.0 (Invitrogen, Cat. No. AM9855G) and 5 µL proteinase K (100 µg; Invitrogen, Cat. No. 25530049), corresponding to a final 1:200 dilution. Samples were incubated for 20 min at 56 °C.

Following PK digestion, 10 µL of the reaction were diluted in 90 µL of dilution buffer (1:2,000 dilution) and incubated at 90 °C for 15 min to inactivate proteinase K. From this dilution, additional serial dilutions were prepared by mixing 10 µL sample with 90 µL dilution buffer to obtain final dilutions of 1:20,000; 1:200,000; 1:2,000,000; and 1:20,000,000. Dilution ranges were selected according to production stage: 1:20,000 for crude cell lysates, 1:200,000–1:2,000,000 following affinity chromatography purification, and 1:2,000,000–1:20,000,000 following iodixanol ultracentrifugation and sample concentration.

The ddPCR reaction mix was prepared with 11 µL of 2× ddPCR Supermix for Probes (No dUTP; Bio-Rad, Cat. No. 1863025), 4.4 µL nuclease-free water, and 1.1 µL of 20× TaqMan primer–probe mix targeting the SV40 promoter (Assay ID: APT2AWV; forward primer: GGAGTGGCACCTTCCA; reverse primer: CTTTATTTGTGAAATTTGTGATGCTATTGCT; probe: GGAACCCCTAGTGATGGAGTT). A volume of 5.5 µL diluted sample was added to each well of a ddPCR semi-skirted 96-well plate (Bio-Rad, Cat. No. 12001925), followed by 16.5 µL of master mix per well. Plates were sealed using a PX1 plate sealer (Bio-Rad Laboratories), vortexed, and briefly centrifuged.

Droplets were generated using a Bio-Rad QX200 automated droplet generator by combining 20 µL droplet generation oil (Bio-Rad, Cat. No. 1863005) with 20 µL reaction mix. PCR amplification was performed using the following cycling conditions: 95 °C for 10 min; 40 cycles of 95 °C for 30 s and 60 °C for 1 min; and a final enzyme deactivation step at 98 °C for 10 min. Droplets were read using a Bio-Rad QX200 droplet reader.

Acceptance criteria for ddPCR data analysis included ≥10,000 accepted droplets, ≥100 negative droplets, and ≥5–10 positive droplets (depending on the negative control). Viral genome titers were calculated as vg/mL and vg/cell as following:

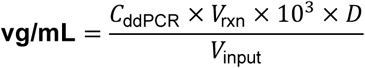

where:

- *C*_ddPCR_= copies per µL measured by ddPCR
- *V*_rxn_= total ddPCR reaction volume (20 µL)
- *D*= dilution factor of the analyzed sample
- *V*_input_= volume of diluted sample added to the reaction (5 µL)
- 10^3^converts µL to mL

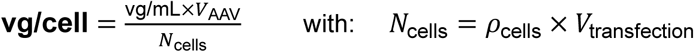

where:

- *V*_AAV_= total volume of AAV sample at the analyzed stage (mL)
- *ρ*_cells_= cell density at the time of transfection (cells/mL)
- *V*_transfection_= transfection culture volume (mL)
- *N*_cells_= total number of cells at transfection

### Agilent 2100 Bioanalyzer based quantification of AAV capsid particles

AAV capsid particle concentration for Figure 3e only was determined indirectly from capsid protein abundance using the Agilent 2100 Bioanalyzer and the Protein 230 Kit according to the manufacturer’s instructions. Affinity-purified AAV samples were analyzed under denaturing reducing conditions. Samples were prepared by mixing 4 μL of AAV material with 2 μL of Protein 230 sample buffer containing 3.5% (*vv*)1 M DTT, followed by heating at 95 °C for 5 min. After brief cooling to room temperature, samples were diluted with 84 μL of deionized water, and 6 μL of the diluted material was loaded onto the Protein 230 chip. The kit ladder was prepared in parallel and used as the sole calibration reference. Chips were prepared using the provided gel-dye mix and destaining solution and runs were initiated within 5 min of sample loading. Separation was performed on the Agilent 2100 Bioanalyzer and data were analyzed using Agilent 2100 Expert software. The molar concentrations of VP1, VP2, and VP3 were quantified from the corresponding electropherogram peaks and summed to obtain the total concentration of capsid protein subunits (*mol*/*mL*). Viral particle concentration (*VP*/*mL*) was calculated by dividing the summed molar concentration by 60, reflecting the 60 capsid protein subunits that constitute one assembled AAV capsid. All samples were analyzed in duplicate.

### Estimated percentage of full capsids based on vg/VP ratio after affinity chromatography purification

The proportion of genome-containing particles in affinity chromatography-purified AAV fractions, Figure 3e only, was estimated using vector genome titers and total particle titers. Vector genomes (*vg*/*mL*) were quantified by ddPCR, whereas total viral particles (*VP*/*mL*) were determined by Agilent 2100 Bioanalyzer analysis of VP1, VP2, and VP3 capsid proteins. The percentage of full capsids was estimated as (*vg*/*mL* ÷ *VP*/*mL*) × 100, based on the assumption that each full capsid contains one packaged vector genome.

### Thermal stability of AAV capsids by nano-differential scanning fluorimetry

The thermal stability of AAV capsids was assessed by nano-differential scanning fluorimetry (nDSF) using a Prometheus PANTA C instrument (NanoTemper Technologies GmbH). AAV samples were analyzed at concentrations ranging from 1.0 × 10¹² to 5.0 × 10¹² vp/mL. Intrinsic protein fluorescence was monitored during a temperature ramp from 20°C to 100°C at a heating rate of 1.0°C/min over a total acquisition time of 80 min. The onset temperature of unfolding (T_on) and melting temperature (T_m) were determined using PANTA Analysis software according to the manufacturer’s guidelines.

Normalized first derivative of 330 nm / 350 nm ratio was used to determine the capsid unfolding temperature (T_M_).

### AAV Genome release by Differential Scanning Fluorimetry (DSF) using SYBR Gold

AAV genome release during capsid unfolding was evaluated using a dye-based DSF assay that monitors the binding of SYBR™ Gold nucleic acid stain (Invitrogen) to exposed or released viral genomes. A total of 3.5 µL of a 143× SYBR Gold working solution was added to 21.5 µL of AAV sample at genomic concentrations ranging from 5.0 × 10¹¹ to 5.0 × 10¹² vg/mL. Samples were sealed, centrifuged for 2 min at 1,000 × g, and subjected to a temperature gradient from 20°C to 100°C at a heating rate of 0.5°C/min (total duration: 160 min) using a Bio-Rad CFX thermocycler. Fluorescence was recorded in the FRET channel (excitation ∼450–490 nm; emission ∼560–580 nm). Normalized fluorescence change (dRFU) was used to determine genome release melting temperatures (T__M_, _DNA_).

### Size-exclusion chromatography with multi-angle light scattering detection (SEC-MALS)

SEC-MALS analyses were performed using a Vanquish UPLC system (Thermo Fisher Scientific) equipped with a diode array detector (DAD). Unless otherwise specified, 2.5 × 10¹¹ vp of AAV sample were injected per run. Protein separation was achieved under isocratic conditions at a flow rate of 0.4 mL/min using a BEH SEC column (450 Å, 2.5 µm, 4.6 × 300 mm; Waters) maintained at 40°C. The mobile phase consisted of 50 mM sodium phosphate (Sigma-Aldrich, Cat. No. 342483), 400 mM sodium perchlorate (Merck, Cat. No. 410241), and 0.01% Pluronic F-68 at pH 6.0. UV absorbance was monitored at 210, 260, and 280 nm. MALS analysis was performed using a DAWN8 MALS detector (Wyatt Technology) coupled to a refractive index (RI) detector (Optilab UT-rEX, Wyatt). System calibration was performed using 99.8% anhydrous toluene (Merck, Cat. No. 244511), and normalization was achieved using bovine serum albumin (BSA), which was also used to correct inter-detector (DAD to MALS and RI) delays. Concentration determination was based on RI measurements using dn/dc values of 0.185 mL/g for protein capsid and 0.170 mL/g for nucleic acids. UV extinction coefficients were 2.1 and 1.3 mL·mg⁻¹·cm⁻¹ at 280 nm and 260 nm for protein, and 14.6 and 24 mL·mg⁻¹·cm⁻¹ at 280 nm and 260 nm for nucleic acids, respectively. The expected molar mass of the AAV capsid protein was 3.75 MDa. Data acquisition and analysis were performed using standard Wyatt software protocols. **Microfluidic capillary gel electrophoresis**

High-throughput microfluidic chip-based capillary gel electrophoresis (HT CE-SDS) analyses of AAV samples were performed using a Caliper LabChip GXII Touch system (PerkinElmer). AAV samples at a concentration of 2.0 × 10¹³ vp/mL were prepared by mixing 20 µL of sample with 35 µL of reducing sample buffer containing 700μl HT Protein Exact buffer supplemented and 12.3μL of 2 M DTT. Samples were heated at 95°C for 5 min, cooled to room temperature, and diluted with 55 µL of water prior to analysis. Samples were analyzed using the HT Protein Exact protocol, and chip preparation was performed according to the manufacturer’s instructions.

### Charge detection mass spectrometry

Charge detection mass spectrometry (CD-MS) was used to assess the percentage of empty, full and overfilled capsids. The measurements were carried out using an electrostatic linear ion trap instrument equipped with an embedded detection cylinder (Megadalton Solutions, Inc.). Individual ion mass-to-charge ratios (m/z) and charge states were measured at an ion energy of 130 eV/z and a trapping time of 104.6 ms, resulting in a charge uncertainty of approximately 0.8 elementary charges. Transient signals were analyzed using frequency chasing. Samples were introduced via nano-electrospray ionization using a Triversa Nanomate (Advion Interchim Scientific). To minimize adduct formation and allow for high resolution mass measurements, 20 µL of each purified AAV sample was buffer exchanged into 200 mM ammonium acetate (ThermoFisher, Cat. No. AM9070G) containing 0.01% Pluronic F-68 (ThermoFisher) using two Micro Bio-Spin P6 columns (Bio-Rad). Clarified lysates were pretreated using a detergent removal column (Pierce, ThermoFisher Scientific) prior to concentration with Vivaspin 500 centrifugal filters (VS0162, Sartorius) followed by buffer exchange into ammonium acetate using Micro Bio-Spin columns. Lysate samples were analyzed using pulsed-mode CD-MS to enhance sensitivity.

### AAV genome integrity assessment by agarose gel electrophoresis

To evaluate the integrity of the encapsidated genetic cargo (single-stranded eGFP cassette, 2,983 bp) across different rAAV serotypes, viral genomes were extracted and analyzed by native and alkaline agarose gel electrophoresis to detect potential genome truncations or fragmentation.

To remove non-encapsidated (free) DNA prior to genome release, a total of 10 µg of rAAV-associated DNA were treated with 2 units (U) of DNase I (Invitrogen, Cat. No. AM2222) diluted in 10× DNase buffer (Invitrogen, Cat. No. AM8170G) and incubated for 30 min at 37 °C. DNase I activity was subsequently terminated by the addition of Tris–EDTA buffer (final EDTA concentration 5 mM, pH 8.0; Merck, Cat. No. T9285). Digested free DNA was removed by filtration using 0.5 mL Amicon centrifugal filters with a 100 kDa molecular weight cut-off (MWCO) (Millipore, Cat. No. UFC505024). Purification of AAV genomic DNA was performed using the QIAquick PCR Purification Kit (Qiagen, Cat. No. PN28104) according to the manufacturer’s instructions. All centrifugation steps were carried out at 17,900 × g at room temperature using a benchtop centrifuge (Eppendorf, Cat. No. 5425). Briefly, five volumes of Buffer PB were added to the DNase-treated samples, followed by incubation at 95 °C for 10 min to release the viral genomes from AAV capsids. Samples were then applied to QIAquick columns and centrifuged for 60 s. After one wash with 750 µL Buffer PE, columns were centrifuged twice for 1 min to remove residual wash buffer, and the purified DNA was eluted in 50 µL Low-TE buffer following a 1 min incubation, in DNA LoBind tubes (Eppendorf, Cat. No. 0030108051). Purified DNA concentrations were determined using the Qubit dsDNA High Sensitivity Assay Kit (Invitrogen, Cat. N° Q32851) according to the manufacturer’s instructions.

For analysis of genomes under native conditions, agarose gel electrophoresis was performed using the Lonza FlashGel™ system (Lonza, Cat. No. 57067) following the manufacturer’s guidelines. A total of 50 ng purified DNA (4 µL, diluted in nuclease-free water if necessary) was mixed with 1 µL FlashGel loading dye and loaded onto the gel. Electrophoresis was conducted at a constant voltage of 250 V for 8 min.

To assess genome integrity under denaturing conditions, alkaline agarose gel electrophoresis was performed. A 1.2% agarose gel was prepared by dissolving two TopVision agarose tablets (Thermo Fisher Scientific, Cat. No. R2901) in 83 mL agarose dissolving buffer (30 mM NaCl (Thermo Fisher, Cat. No. A57006), 2 mM EDTA, pH 7.5) by heating in a microwave until fully dissolved. The molten agarose was cast into a gel cassette fitted with 20-well single combs and allowed to solidify at 4 °C for 30 min. The gel was then equilibrated overnight at 4 °C in alkaline running buffer containing 30 mM NaOH (Sigma-Aldrich, Cat. No. 4.80648.2500) and 2 mM EDTA (pH 12.5). For sample preparation, 150 ng purified DNA diluted in nuclease-free water (16.7 µL total volume) were mixed with 3.34 µL of 6× alkaline agarose gel loading dye (Thermo Fisher Scientific, Cat. No. J62157-AB) and heated at 90 °C for 5 min. Samples (20 µL) and 10 µL of E-Gel™ 1 kb DNA ladder (Thermo Fisher Scientific, Cat. No. 10488090) were loaded into the pre-cooled alkaline gel and electrophoresed at a constant voltage of 100 V for 85 min at 4 °C. Following electrophoresis, the gel was rinsed twice with neutralization buffer (0.5 M Tris-HCl, pH 7.5; Thermo Fisher Scientific, Cat. No. 15567027) and incubated for 30 min in the same buffer. DNA was visualized by staining the gel for 1 h at room temperature in the dark with 10 µL SYBR™ Gold nucleic acid stain (Thermo Fisher Scientific, Cat. No. S11494) prepared in 100 mL of 1× TBE solution (Thermo Fisher Scientific, Cat. No. 28355). Gels were rinsed once with distilled water and imaged using a ChemiDoc™ MP Imaging System (Bio-Rad, Cat. No. 12003154). Genome integrity was defined by the presence of a predominant band corresponding to the expected 2,983 bp size.

### Cell culture

In the current study, multiple cell types of cells were used to assess and compare the *in vitro* transduction efficiency of different distant and human/NHP-derived AAVs.

#### Human Embryonic Kidney (HEK-293T) cells

(Thermo Fisher Scientific) were maintained in suspension culture in FreeStyle Media 293 containing Glutamax supplement (Gibco, Cat. No. 12338018) at 2.00E+05 viable cells (vc)/mL (37°C, 6% CO_2_, 110 RPM). One day prior to transduction assay, cells were split and cultured at a density of 5.00E+05 vc/mL to promote a cell growth phase. For transduction assays, cells were seeded into black, optically clear bottom, cell culture-treated 96 or 384 well plates - (Thermo Scientific, 96 well - Cat. No. 165305, 384 well – Cat. No. 142761). After allowing 1–2 h for cell attachment, AAV vectors diluted in FreeStyle™ Medium were added on the same day.

#### Primary human lung small airway epithelial cells (SAEC)

(Lifeline Cell Technologies, FC-0106, Lot 08363) were thawed in BronchiaLife Epithelial Airway Complete Medium (Lifeline Cell Technologies, Cat. No. LL-0023) containing basal medium, 6mM L-Glutamine, TM1 Life Factors (1uM Epinephrine, 5ug/mL Transferin PS, 10nM T3, 0.1ug/mL Hydrocortisone, 5ng/mL rh EGF, 5ug/mL rh Insulin), 0.4% Extract P and HLL supplement (500ug/mL HSA, 0.6uM Linoleic Acid, 0.6 ug/mL Lecithin).

#### Primary human bronchial/ tracheal epithelial cells

(ATCC, PCS-300-010, Lot 70035985) were thawed in airway epithelial cell basal medium (ATCC, Cat. No. PCS-300-030) supplemented with 6mM L-Glutamine, 1mM Epinephrine, 5mg/mL Transferin, 10nM T3, 5mg/mL Hydrocortisone, 5ng/mL rh EGF, 5mg/mL rh Insulin), 0.4% Extract P and HLL supplement (500mg/mL HSA, 0.6mM Linoleic Acid, 0.6 mg/mL Lecithin) (ATCC, Cat. No. PCS-300-040).

#### Human lung microvascular endothelial cells

(Innoprot, P10552, Lot 325K272) were thawed in complete microvascular endothelial cell medium (Innoprot, Cat. No. P60104) containing endothelial cell basal medium, 5% FBS, 1% endothelial cell growth supplement and 1% penicillin/ streptomycin (P/S) solution. Their affiliated culture wells were coated with 2ug/cm^2^ fibronectin diluted in sterile water (Innoprot, Cat. No. P8248) at 37°C for 1-2h before cell plating.

All adherent cell lines used in this study were thawed and directly plated in 20 mL complete growth medium in T75 cm² cell culture–treated flasks (Greiner Bio-One, Cat. No. 658175) at a density of 4.00 – 5.00E+05 cells per flask. Cells were expanded under standard culture conditions until reaching 70–80% confluency. For passaging and plating the cells for transduction assay, cells were washed once with 1× DPBS and detached by incubation with 3 mL TrypLE™ Express cell dissociation enzyme (Gibco, Cat. No. 12604-013) at 37 °C for 3–5 min. Enzymatic dissociation was stopped using 1× DPBS supplemented with 5% fetal bovine serum (FBS), followed by centrifugation for 5 min at 250 × g (A549 cells), 120 × g (COS-1 cells), or 150 × g (small airway epithelial cells, bronchial/tracheal epithelial cells, and cardiac microvascular endothelial cells). Cells were then resuspended in their corresponding complete medium and seeded into black, optically clear-bottom, cell culture–treated 96-or 384-well plates at densities recommended by the respective manufacturers (Table S4). Twenty-four hours after plating, allowing for cell attachment and recovery, AAV vectors were added to the cultures and transduction efficiency was assessed 72 h later.

#### Primary human pulmonary alveolar epithelial cells

(Innoprot, P10556, Lot 425K031) were thawed in complete alveolar epithelial cell medium (Innoprot, Cat. No. P60102) containing 2% FBS, 1% epithelial cell growth supplements and 1% P/S solution. Their affiliated culture wells were coated with 2ug/cm^2^ poly-L-lysine diluted in sterile water (Innoprot, Cat. No. PLL) at 37°C for 1-2h before cell plating. Considering that these cells do not proliferate in culture, in contrast to the other cell types, AAVs were added to them 2h after seeding and transduction was assessed 72h later.

### AAV Transduction assay

To evaluate the infectivity of distant or human/non-human primate (NHP)–derived recombinant AAVs across multiple cell types, vectors were diluted in the corresponding complete culture medium for each cell type, without fetal bovine serum (FBS). Four multiplicities of infection (MOIs) were tested (1.0 × 10⁶, 5.0 × 10⁵, 2.5 × 10⁵, and 1.25 × 10⁵ vg/cell). Except for HEK293T cells, which were seeded and transduced on the same day, and primary mouse cortical neurons, which were plated 8 days prior to transduction, all other cell types were seeded 24 h before AAV addition. On the day of transduction, culture medium was removed and adherent cells were washed once with 1× DPBS. AAV preparations diluted to the indicated MOIs were then added to the cells and incubated for 3 h at 37°C and 5% CO₂. For cell types requiring FBS supplementation, the appropriate concentration of FBS was added after the initial 3 h incubation to minimize serum-mediated interference with viral entry. Cells were subsequently maintained for 72 h at 37°C and 5% CO₂ before eGFP transgene expression was assessed by fluorescence microscopy and flow cytometry.

The same transduction protocol was used to determine the MOI required to transduce 80–90% of HEK293T cells, a parameter necessary for subsequent AAV neutralization assays. For this purpose, 11 serial MOIs were tested in a 1:2 dilution series, ranging from 1.0 × 10⁶ to 9.77 × 10² vg/cell.

### Detection and quantification of GFP positive cells by fluorescence microscopy and flow cytometry

GFP expression was first assessed qualitatively by fluorescence microscopy using a ZOE Fluorescent Cell Imager (Bio-Rad Laboratories, Cat. No. 1450031). After visual inspection of all four multiplicities of infection (MOIs) for cell health and viability, images were acquired for the 1.0 × 10⁶ vg/cell condition from three independent replicates, with three distinct fields of view (FOVs) per replicate, using the GFP channel (excitation 488 nm, emission 500–550 nm; scale bar: 100 µm).

Quantitative analysis of GFP-positive cells was performed by flow cytometry. Cells were washed with 1× DPBS and detached using TrypLE Express cell dissociation enzyme (Gibco, Cat. No. 12604-013). Enzymatic detachment was stopped by the addition of 1× DPBS supplemented with 10% fetal bovine serum (FBS), followed by centrifugation. Cell pellets were resuspended in FACS buffer consisting of 1× DPBS, 2% heat-inactivated fetal calf serum (Cytiva Life Sciences, Cat. No. SH30073.03HI), and 2 mM EDTA. Cell suspensions (40 µL for 384-well plates or 60 µL for 96-well plates) were acquired on a BD FACSymphony A5 SE flow cytometer (BD Biosciences) using the blue 510 nm detection channel for eGFP. For each MOI condition, three biological replicates were analyzed, and untransduced cells were included as negative controls.

Flow cytometry data were analyzed using FlowJo software (v10.10.0). Viable cells were identified based on forward and side scatter (FSC-A vs SSC-A), and singlets were gated using FSC-H vs FSC-A. The GFP-positive gate was defined using untransduced control samples, and both the percentage of GFP-positive cells and the median fluorescence intensity (MFI) were determined. Mean values and standard deviations were calculated from the two / three replicates (depending on cell number availability).

### Enzyme-linked immunosorbent assay (ELISA)

Enzyme-linked immunosorbent assay (ELISA) was used to quantify total anti-AAV binding antibodies in mouse and human serum samples. Total anti-AAV IgG and IgM antibodies were measured in mouse sera collected from two independent in vivo studies (Supplemental Figs. 5 and 6), while total anti-AAV IgG antibodies were assessed in human sera.

Recombinant AAV capsids (AAV1, AAV2, AAV4, AAV5, AAV8, AAV9, and Bovine AAV) were diluted in 0.1 M carbonate–bicarbonate coating buffer (Thermo Scientific, Cat. No. 28382) to a final concentration of 1.0 × 10¹⁰ viral particles (vp)/mL. Fifteen microliters of each diluted AAV preparation were dispensed into clear, flat-bottom immuno 384-well plates (Thermo Scientific, Cat. No. 464718) using an electronic 384-channel pipette (Integra ViaFlow 384, Cat. No. 6031) and plates were incubated overnight at +4°C. Control plates containing carbonate-bicarbonate buffer without any AAV were included as negative control.

Following coating, plates were washed twice with 100 µL wash buffer consisting of 1× DPBS (Gibco, Cat. No. 14190144) supplemented with 0.025% Tween-20 (Sigma-Aldrich, Cat. No. P1379) using a BioTek BioStack microplate stacker (Agilent). Plates were then blocked with 40 µL blocking buffer (1× DPBS containing 0.5% BSA; Sigma-Aldrich, Cat. No. A7906) for 1 h at room temperature or overnight at 4 °C to prevent non-specific binding.

Serum samples were diluted in blocking buffer, starting with an initial 1:50 dilution followed by ten additional serial dilutions at a 1:3 ratio, each prepared from the preceding dilution. After removal of the blocking buffer, 15 µL of each serum dilution were added to the coated plates and incubated overnight at 4 °C. The following day, plates were washed twice with wash buffer and incubated with 15 µL of horseradish peroxidase (HRP)-conjugated secondary antibodies diluted 1:5,000 in blocking buffer. The following secondary antibodies were used as appropriate: goat anti-mouse IgM-HRP (SouthernBiotech, Cat. No. 1021-05) and goat anti-mouse IgG-HRP (SouthernBiotech, Cat. No. 1036-05). Plates were incubated for 1 h at room temperature or overnight at 4 °C.

After two additional washes with wash buffer, plates were developed by adding 15 µL of tetramethylbenzidine (TMB) substrate solution (SeramunBlau fast2, Seramun Diagnostics GmbH, Cat. No. S-100-TMB) and incubated at room temperature until positive control wells developed a visible signal (typically 3–5 min). The reaction was stopped by addition of 15 µL of 250 mM sulfuric acid (Honeywell, Fluka), and absorbance at 450 nm was measured immediately using a Synergy H1 microplate reader (BioTek, Agilent). Antibody titers were defined as the highest serum dilution yielding an absorbance value greater than two-fold above the mean absorbance of 15 negative-control wells lacking serum.

### Neutralization / Transduction inhibition cellular assay

To assess the neutralizing activity of anti-AAV antibodies present in sera from AAV-immunized mice or healthy human donors, a cell-based transduction inhibition assay was performed using HEK293T cells. Cells were split one day prior to the assay to ensure active proliferation at the time of infection. On the day of the experiment, 3,200 cells per well were seeded in black, optically clear-bottom, cell culture-treated 384-well plates (Thermo Scientific, Cat. No. 142761) in FreeStyle™ 293 Expression Medium supplemented with GlutaMAX™ (Gibco, Cat. No. 12338018) and allowed to attach for 1–3 h at 37°C and 5% CO₂.

In parallel, recombinant AAV vectors were diluted in FreeStyle 293 medium at serotype-specific concentrations previously determined to achieve 80–90% transduction of HEK293T cells in a standard transduction assay. The final vector doses used were: AAV1 (1.25 × 10⁵ vg/cell), AAV2-ESGHGYF (5.00 × 10⁵ vg/cell), AAV2 (4.50 × 10³ vg/cell), AAV4 (8.00 × 10⁵ vg/cell), AAV5 (1.00 × 10⁶ vg/cell), AAV8 (1.00 × 10⁶ vg/cell), AAV9 (8.00 × 10⁵ vg/cell), and bovine AAV (1.00 × 10⁵ vg/cell).

Serum samples were diluted in FreeStyle 293 medium to an initial final assay dilution of 1:20, followed by four serial 1:4 dilutions prepared from the preceding dilution. Diluted sera were incubated with the corresponding rAAV preparations for 1 h at 37°C to allow antibody–capsid binding. The serum–virus mixtures were then added to the cells. After 3 h of incubation, fetal bovine serum (FBS) was added to a final concentration of 8.3% to minimize serum-mediated interference with viral entry. Cells were subsequently maintained for 72 h at 37°C and 5% CO₂.

Transgene expression was assessed by flow cytometry based on the percentage of GFP+ cells detected, as described above. Cells incubated with rAAVs in the absence of serum served as positive (no-neutralization) controls, whereas cells incubated with medium alone (no rAAV, no serum) served as negative controls. When sufficient serum volume was available, samples were analyzed in duplicate and the mean value was reported. Sera were considered neutralizing if a 1:20 dilution blocked at least 50% of vector transduction.

Percent neutralization was calculated as:

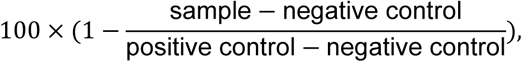

where the positive control represents fully transduced cells and the negative control represents untransduced cells.

### *In vivo* animal experiments

All animal experiments performed in this study were approved by and conducted in accordance with the guidelines of the Swiss Cantonal Veterinary Office (Basel) and the Novartis Animal Welfare Department.

### Housing and husbandry conditions

Male C57BL/6J mice aged 9–11 weeks were obtained from Charles River Laboratories (Germany). Animals were housed in Allentown XJ individually ventilated cages equipped with automated stainless-steel watering systems (Avidity) and BK 8–15 spruce granulated bedding (J. Rettenmaier & Söhne GmbH + Co. KG). Environmental enrichment consisted of a wooden popsicle stick and wood block (Pura Aspen Chew Sticks and Blocks, Labodia), ≥11 g crinkle paper nesting material (pressed paper strips; J. Rettenmaier & Söhne GmbH + Co. KG), a red polycarbonate mouse house (Tecniplast), and a transparent handling tunnel (Zoonlab).

Animals were provided ad libitum access to gamma-irradiated, vitamin-fortified pelleted diet (Kliba Nafag, Cat. No. 3802) and reverse-osmosis water (hyper-chlorinated, 2–3 ppm). Mice were maintained under a 12 h light/12 h dark cycle (lights on at 06:00, off at 18:00), at a controlled ambient temperature of 22 °C (±2 K) and relative humidity of 55% (range 45–65%). Cages were fully changed every two weeks, with transfer of unsoiled nesting material during cage changes. All animal handling was performed using tunnels to minimize stress.

### Intravenous AAV administration

All recombinant AAV vectors, administered either individually or as pooled preparations, were formulated in buffer containing 20 mM Tris, 1 mM MgCl₂, 200 mM NaCl, and 0.005% Pluronic (pH 8.1). Prior to injection, mice were placed in a warming device to promote tail vein dilation. Intravenous injections were performed via the lateral tail vein using a total injection volume of 100–120 µL, corresponding to approximately 4 mL/kg body weight. Following injection, gentle pressure was applied to the injection site to prevent bleeding, and animals were monitored until full recovery. Post-injection, animal well-being was monitored throughout the study duration. Clinical observations included body condition (weight loss, injection site reactions, ruffled fur, wet or soiled perianal region), posture (normal vs. hunched), behavior (motility, nesting activity, signs of apathy), and respiration (normal vs. labored or irregular).

Of note, for comparison of vector transduction in the presence or absence of prior AAV pool immunization, the experiments presented in Figures 1 and 3 were performed in parallel using mice from the same cohort which were matched for age and sex and processed under identical experimental conditions. Figure 1 includes the non-immunized control (No AAV) and vector-treated groups (AAV2-ESGHGYF-GFP / AAV4-GFP / AAV9-GFP and Bovine AAV-GFP), whereas Figure 3 includes the corresponding AAV pool (empty capsids)-immunized groups challenged with no vector, AAV4-GFP, AAV9-GFP, or Bovine AAV-GFP. The non-immunized groups (-AAV pool) were therefore used both for the analyses presented in Figure 1 and as reference comparators in Figure 3. This study design enabled direct cross-figure comparison while reducing the total number of animals used.

### Vena saphena blood collection

Blood samples were collected from the lateral saphenous vein without anesthesia. Mice were gently restrained, the hind limb was extended, and the collection site was shaved and disinfected with 70% ethanol. The vein was punctured using a 23 G needle, and approximately 70 µL of blood was collected into CB300 Serum CAT Microvette tubes (Sarstedt, Cat. No. 16.440). Hemostasis was achieved by applying gentle pressure with sterile gauze. Collected blood was allowed to clot for 20 min at room temperature, followed by centrifugation at 2,500 × g for 10 min. Serum (approximately 25–30 µL) was carefully transferred into sterile protein low-binding tubes (Eppendorf, Cat. No. 0030108094) and stored at −80 °C until further analysis.

### Mouse euthanasia and organ collection

At the experimental endpoints (28 or 49 days post-AAV administration, depending on study objectives), mice were deeply anesthetized and positioned in the supine position. The thoracic cavity was opened to expose the heart. When required, terminal blood samples were collected from the right ventricle using S-Monovette Z 1.2 mL tubes (Sarstedt, Cat. No. 06.1663.001). A cannula was inserted into the left ventricle, and the right atrium was incised to allow efflux. Transcardiac perfusion was performed using 1× DPBS for 3 min at a flow rate of 5 mL/min to remove circulating blood. Following perfusion, organs including the lungs, liver, brain, heart, and kidneys were harvested. From each organ, a representative tissue sample (approximately 50–100 mg) was placed in RNAlater stabilization solution (Invitrogen, Cat. No. AM7022) and stored at −80 °C for downstream molecular analyses. The remaining tissue was processed for immunohistochemical assessment of AAV biodistribution at the cellular level, as described below.

### Animal tissue processing and immunohistochemistry

All collected tissues were fixed in 10% neutral buffered formalin for 48 h at room temperature, followed by trimming and paraffin embedding according to standard procedures. Formalin-fixed, paraffin-embedded (FFPE) tissue blocks were sectioned at 3 µm thickness, and sections were mounted onto SuperFrost® Plus slides (Thermo Fisher Scientific, Switzerland) for immunohistochemical analysis.

Immunohistochemical stainings were performed using automated Ventana instruments (Discovery® ULTRA; Roche Diagnostics Schweiz AG, Rotkreuz, Switzerland). Tissue sections were deparaffinized and rehydrated under solvent-free conditions, followed by heat-induced antigen retrieval. Sections were incubated for 1 h at room temperature with a rabbit polyclonal anti-GFP antibody (Thermo Fisher Scientific; Cat. No. A-11122) diluted 1:1,000 in antibody diluent. Signal detection was achieved using an anti-rabbit HRP-conjugated multimer secondary antibody (OmniMap® anti-Rabbit HRP, ready-to-use; Roche, Cat. No. 05269679001) and 3,3′-diaminobenzidine (DAB) chromogen (ChromoMap® kit; Roche, Cat. No. 05266645001), according to the manufacturer’s instructions. Slides were counterstained with hematoxylin and bluing reagent, dehydrated, and mounted using Eukitt® mounting medium (Medite, Dietikon, Switzerland).

Whole-slide digital images were acquired using a NanoZoomer S60 slide scanner (Hamamatsu Photonics; NDP-Scan software v2.5). Staining patterns were evaluated by visual examination of the digital scans.

### Immunofluorescence staining of FFPE-lung sections

To determine the cellular identity of transduced cells in the lung, double immunofluorescence stainings were performed on 3μm FFPE lung sections. All stainings were performed using the automated Discovery® ULTRA instrument, using reagents from Roche Diagnostics and conditions similar to the immunohistochemistry procedure described above. Vector-transduced cells were detected using chicken anti-GFP (Abcam Ltd., Cat. No. ab13970) diluted at 1:500 in Ventana antibody diluent (Roche, Cat. No. 06440002001). For cell-type identification, sections were co-stained with rabbit anti-aquaporin 5 (AQP5, Abcam, Cat. No. ab 315655), diluted at 1:20,000 to label alveolar type I pneumocytes, rabbit anti-surfactant protein C (SFTPC, Abcam, Cat. No. ab322443, clone RM1264), diluted at 1:500 to label alveolar type II pneumocytes, or rat anti-CD31 (Dianova: Cat. No. DIA310), diluted at 1:20 to label pulmonary endothelial cells. All primary antibodies were incubated for 3 hours at room temperature. Detection of the vector-transduced cells was performed with an anti-chicken secondary antibody (Donkey Anti-Chicken HRP, Cat. No. 703-036-155, Jackson Immunoresearch, diluted at 1:200 in Ventana diluent) followed by the Discovery FITC Kit (Cat. No. 07259212001). The cell type markers were detected with either an anti-rabbit (ready to use OmniMap® anti-Rabbit HRP, Cat. No. 05269679001) or an anti-mouse (ready to use OmniMap® anti-Rat HRP, Cat. No. 05891892001) and the Discovery Rhodamine Kit (Cat. No. 07259883001) following the provider’s recommendations. Nuclei were counterstained with the Discovery QD DAPI kit (Cat. No. 05268826001). Fluorescent images were acquired by confocal microscopy using an Olympus confocal microscope (model FV3000) equipped with a 60x objective to assess co-localization of GFP-positive cells with the indicated lung cell-type markers.

### Genomic DNA & total RNA isolation from harvested tissue

Genomic DNA and total RNA were simultaneously isolated from harvested tissues using the AllPrep DNA/RNA kit according to the manufacturer’s instructions (Qiagen, Cat. No. 80311). Briefly, 5–10 mg of tissue were transferred into 2 mL screw-cap tubes containing 2.8 mm ceramic–zirconium oxide beads (Precellys, Cat. No. P000916-LYSK0-A). Tissues were lysed in 350 µL RLT lysis buffer supplemented with β-mercaptoethanol (3.5 µL per sample; Merck, Cat. No. 60-24-2) and homogenized using a Precellys 24 Touch homogenizer (Bertin Technologies) at 5,500 rpm for 20–24 s. Homogenized lysates were stored at −80 °C until further processing.

On the day of nucleic acid extraction, samples were thawed at room temperature (20 °C), and all subsequent steps, including centrifugations, were performed at this temperature. Lysates were centrifuged at 5,600 × g for 4 min to pellet insoluble debris. A total of 300 µL of the clarified supernatant was transferred to an AllPrep 96-well DNA plate placed on top of an S-Block. The plate was sealed and centrifuged at 5,600 × g for 4 min. Following centrifugation, the DNA-binding plate was sealed and stored at +4 °C for subsequent DNA purification, while the flow-through containing total RNA was processed immediately.

### RNA isolation

For RNA isolation, 300 µL of the RNA-containing flow-through was mixed with an equal volume (300 µL) of 70% RNase-free ethanol. The mixture was transferred to an RNeasy 96-well plate placed on a clean S-Block, sealed with Airpore tape, and centrifuged at 5,600 × g for 4 min to allow RNA binding to the membrane. Bound RNA was washed once with 800 µL Buffer RW1 and twice with 800 µL Buffer RPE, with centrifugation steps performed between washes. RNA was eluted into Elution Microtubes CL using a total of 100 µL RNase-free water, applied in two sequential steps (50 µL + 50 µL) with a 2 min incubation between elutions.

### DNA isolation

The AllPrep 96-well DNA plate containing bound genomic DNA was washed sequentially with 800 µL Buffer AW1 and 800 µL Buffer AW2, with centrifugation steps performed between washes. Genomic DNA was eluted using 100 µL Buffer EB pre-warmed to 70 °C, applied in two steps (50 µL + 50 µL) with a 5 min incubation between elutions.

Purified DNA and RNA samples were stored at −80 °C until further analysis.

### RNA/ DNA concentration quantification

DNA and RNA concentrations were quantified using a Qubit Fluorometer with DNA and RNA Broad Range (BR) assays, according to the manufacturer’s instructions (Invitrogen; DNA BR Cat. No. Q32853; RNA BR Cat. No. Q10211). Briefly, 1 µL of each DNA or RNA sample was mixed with 199 µL of the respective BR working solution, briefly vortexed, incubated for 2 min at room temperature, and measured using a GloMax Discover Reader (Promega, Cat. No. GM3000). Triplicates of low and high reference standards were measured in parallel, and sample concentrations were normalized to the average values obtained for the corresponding standards.

### Quantification of AAV biodistribution at the DNA level in tissue samples

AAV biodistribution in tissue samples was quantified as vector genomes per diploid genome (vg/dg) using droplet digital PCR (ddPCR). An AAV-specific primer–probe set targeting the SV40 polyadenylation signal (SV40pA133) was used to quantify vector genomes, whereas the murine *Rpp30* gene, present as a single copy per haploid genome, was used as an internal reference to determine the number of host diploid genomes in each DNA sample.

Prior to ddPCR, concentrated genomic DNA samples were diluted in nuclease-free water to a final concentration of 5 ng/µL. The ddPCR reaction master mix was prepared as follows: 1.1 µL AAV-specific FAM-labeled primer–probe mix targeting the SV40pA133 late (3′-end) polyadenylation signal (forward primer: TGCTTTATTTGTGAAATTTGTGATGCT; reverse primer: CCCTGAACCTGAAACATAAAATGA; probe: TGTAACCATTATAAGCTGCAATAAACAAGTTAACAACAACA), 1.1 µL Rpp30 HEX-labeled primer–probe mix (Bio-Rad Laboratories; assay ID: dMmuCNS822293939), 1.1 µL SphI-HF restriction enzyme mix for genomic DNA digestion (New England Biolabs, Cat. No. R3182S), 11 µL ddPCR Supermix for Probes (No dUTP) (Bio-Rad, Cat. No. 1863024), and 2.2 µL nuclease-free water.

For each reaction, 5.5 µL of diluted DNA sample (corresponding to 20 ng total genomic DNA) was transferred to a ddPCR semi-skirted 96-well plate (Bio-Rad, Cat. No. 12001925) and mixed with 16.5 µL of the prepared master mix. Plates were sealed with a pierceable heat-seal foil using a PX1 plate sealer (Bio-Rad), vortexed briefly, and centrifuged. The final PCR mix was incubated at room temperature for 15 min to allow efficient restriction enzyme digestion of the genomic DNA prior to droplet generation and amplification.

Droplets were generated using a Bio-Rad QX200 Automated Droplet Generator by combining 20 µL of reaction mix with 20 µL of droplet generation oil (Bio-Rad, Cat. No. 1863005). PCR amplification was performed on a C1000 Touch Thermal Cycler (Bio-Rad) equipped with a heated lid (105 °C), using the following cycling conditions: 95 °C for 10 min; 40 cycles of 95 °C for 30 s and 60 °C for 1 min; and a final enzyme deactivation step at 98 °C for 10 min.

Following amplification, droplets were analyzed using a Bio-Rad QX200 Droplet Reader, and absolute copy numbers were determined using QuantaSoft Analysis Pro software (Bio-Rad). AAV biodistribution was expressed as vector genomes per diploid genome (vg/dg), calculated as:

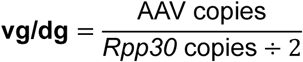

This normalization accounts for variations in DNA input, extraction efficiency, and tissue cellularity, enabling direct comparison of AAV distribution across different tissues.

### Quantification of AAV transgene expression in tissue samples

AAV transgene expression was quantified by reverse transcription (RT)-ddPCR using a primer–probe set targeting the HPRE sequence located in the 3′ UTR of the vector transcript. HPRE provides a highly specific target, absent from endogenous mammalian RNA, and included in all correctly processed AAV-derived transcripts, enabling sensitive, transgene-independent quantification of AAV RNA across tissues.

Because total RNA yield and expected AAV transgene expression varied substantially between tissues, the amount of RNA loaded per RT-ddPCR reaction was adjusted to remain within the assay’s linear dynamic range while maintaining optimal droplet quality. Based on preliminary optimization experiments, the following RNA inputs were used: lungs, 10 ng; liver, 0.1 ng; brain, 50 ng; kidneys, 100 ng; heart, 80 ng. To eliminate residual genomic or vector DNA, RNA samples were treated with 2.73 U DNase I (Qiagen, Cat. No. 79256) for 30 min at room temperature, followed by heat inactivation at 75 °C for 10 min.

RT-ddPCR reactions were assembled on ice using the One-Step RT-ddPCR Advanced Kit for Probes (Bio-Rad, Cat. No. 1864022). Each 22 µL reaction contained 5.5 µL Supermix, 2.2 µL reverse transcriptase, 0.55 µL 300 mM DTT, 1.1 µL HPRE-specific primer–probe mix (forward primer: *TCCATACTGCGGAACTCCTA*; reverse primer: *CAGCCCTGGAAACGATGTAT*; probe: 5’FAM-CAGCAGGTCTGGAGCGAAACTCAT-BHQ1-3’), 6.6 µL nuclease-free water, and the DNase-treated RNA (0.1–100 ng depending on tissue and AAV dose injected).

Droplets were generated using the QX200 AutoDG system (Bio-Rad) with DG32 cartridges and AutoDG oil according to the manufacturer’s instructions. Each droplet consisted of 20µl droplet generation oil and 20µl sample. Thermal cycling was performed on a C1000 Touch Thermal Cycler (Bio-Rad) with a heated lid (105°C) using the following conditions: 50°C for 60 min (reverse transcription), 95°C for 10 min (enzyme activation), followed by 40 cycles of 95°C for 30 s and 55°C for 1 min (denaturation/annealing; 2°C/s ramp rate), and a final 98°C for 10 min enzyme deactivation step.

Droplets were read on a QX200 Droplet Reader, and absolute HPRE transcript counts were determined using QuantaSoft Analysis Pro software (Bio-Rad). Transcript abundance was normalized to the amount of RNA loaded into each reaction and expressed as AAV-RNA copies per µg total RNA, calculated as:

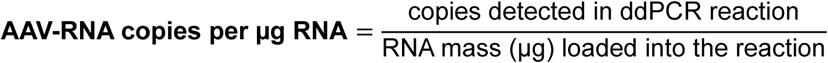

Normalization to RNA mass rather than a reference gene was chosen to avoid variability due to tissue-specific differences in housekeeping gene expression.

## Graphical abstract

The graphical abstract was designed using BioRender software.

## Data and materials availability

All data supporting the results of this study are present within the paper or the Supplemental Materials section.

## Acknowledgements

We thank our collaborators and acknowledge the following contributions to the study: Lisa Trapp, Paulina Baczyk and Jasmin Widmer for their help with the in-depth characterization of AAVs via SEC-MALs, HT CE-SDS and DSF analysis; Louis Fischer, Berangere Gapp, Valerie Cordier, Sabrina Surber and Allen Abey for their support with the *in vivo* mouse experiments; Salvatore Caruso for help with testing AAV productions at small scale; Fabrizio Pizzo, Ralph Riedl and Attila Regös for their help with the automatized ELISA for the characterization of anti-AAV antibodies; Edwige B. Louka for mass-spectrometry analysis of different AAV serotypes, Nathalie Loll, Brigitte Christen, Anna Hudek and Jeannine Hehlen for help with primary mouse spleen cell isolation; Jan Schmidt, Tina Rubic-Schneider and Maria Vono for providing feedback and intellectual input; Benoit Fischer and Francesca Moretti for providing primary human hepatocytes and multiple cell lines for assessment of AAV transduction potential; Erica Coratella and Rebeca Bohnert for guidance during the transduction of primary human hepatocytes; Flavio Schwald and Christelle Gerard for help with confocal microscopy imaging; Sarah Egger, Marie-Christine Lasbennes and Rose-Marie Ungricht for help with testing of Bovine AAV transduction potential on human iPCS-derived kidney organoids; Andreas Hein and Loren Miraglia for intellectual input on investigations related to Bovine AAV ligand; Juliane Kuklik, Claire Domenger and Mirjam Buchs for feedback and intellectual input. Last but not list, we thank the donors for their samples donated to research.

## Author contributions

Project Conceptualization was performed by A.C., A.K., D.C.I., J.M.C. and J.W. Methodology design, establishment and execution involved A.C., A.K., B.T., D.C.I., D.U., E.B., E.E., F.D.J., J.M.C., K.D., L.B., L.I., M.J., M.S., N.C., N.O.A., N.S., S.M., V.C. and V.D. Data analysis and visualization was conducted by C.D., D.C.I., D.U., E.B., J.W., V.D. and K.S. Project supervision included A.C., A.K., D.C.I., J.M.C., J.W., L.B., L.I., V.C. and V.D. Funding was obtained by A.C., D.B., E.T. and J.M.C. Original draft of the manuscript was written by D.C.I. Reviewing, editing and critical evaluation of the content and final version of the manuscript were conducted by all authors

## Declaration of Generative AI and AI-assisted technologies in the writing process

During the preparation of this work the authors used Copilot and Internal ChatGPT to improve readability and language through rephrasing of certain paragraphs. After using these tools, the authors reviewed and edited the content as needed and take full responsibility for the content of the publication.

## Supplemental Figures

Supplemental Figure 1. Phylogenetic tree analysis & similarity at VP1 amino acid level between evolutionarily distant & human/ non-human primate-derived AAVs

Phylogenetic tree featuring the genetic distances between 25 parvoviruses (PV) from eight animal species (avian-AAAV.DA-1, bovine - BAAV, bearded dragon - BDPV, bat - BtAAV, Californian sea lion – Csl-AAV1, duck – DPV, DPV-G018-19, goose – GAAV, and snake - SAAV) (highlighted in blue) alongside 15 human / primate-derived isolates (AAV1, AAV2, AAV3, AAV3b, AAV4, AAV5, AAV6, AAV7, AAV8, AAV9, AAV10, AAV11, AAV12, AAVrh10, AAVrh.74).

The VP1 amino acid sequences of each PV were aligned using the MUSCLE algorithm and this alignment was used to construct the phylogenetic tree. The tree inference was performed using the neighbor-joining method. All analyses were conducted in Geneious Prime 2025 software.

**Supplemental Figure 2. I*n vitro* transduction of primary human hepatocytes transduced with AAV2, AAV2-ESGHGYF, AAV4, AAV6, AAV9, Bovine AAV or Control buffer (No AAV).**

**a) Flow cytometry gating strategy for assessment of primary human lung and liver cells transduction by different AAVs.** 2000 to 10.000 events were acquired using a BD FACSymphony A5 SE flow cytometer. Viable cells were gated based on FSC-A versus SSC-A, followed by singlet gating using FSC-H versus FSC-A. The GFP positive gate was defined using untransduced control cells (No AAV). Representative plots from primary human lung microvascular endothelial cells transduced with No AAV or Bovine AAV are shown.

**b) Overlaid flow cytometry representative histograms** showing GFP fluorescence in primary human lung cell types transduced with AAV2, AAV2-ESGHGYF, AAV4, AAV9, Bovine AAV or buffer control (No AAV) at 1.0 × 10^6^vg per cell.

**c) Representative fluorescence microscopy images** of primary human hepatocytes transduced with AAV2, AAV2-ESGHGYF, AAV4, AAV6, AAV9, Bovine AAV or buffer control 3 days after transduction at MOI 1.0 × 10^6^ vg per cell. Scale bars, 100 μm.

**d) Overlaid flow cytometry representative histograms** showing GFP fluorescence in primary human hepatocytes 3 days after transduction with AAV2, AAV2-ESGHGYF, AAV4, AAV6, AAV9, Bovine AAV or buffer control at MOI 1.0 × 10^6^vg per cell.

**e) Percentage of GFP positive cells** and **f) Median fluorescence intensity of GFP positive cells** quantified by flow cytometry across four tested MOIs (1.0 × 10^6^,5.0 × 10^5^,2.5 × 10^5^,and 1.25 × 10^5^ vg per cell) 3 days after transduction. Data are shown as mean (SD) of 2 technical replicates. Statistical analysis was performed using two-way ANOVA with Dunnett multiple-comparisons test, with each AAV compared to AAV6. Significance is shown relative to AAV6 (∗p<0.05, ∗∗p<0.01, ∗∗∗p<0.001, ∗∗∗∗p<0.0001).

Supplemental Figure 3. Large-scale production, purification, and biophysical characterization of AAV9 and Bovine AAV.

Recombinant AAV9 and Bovine AAV vectors carrying an ss-CAG-GFP genome were produced at large scale in 2.8 liters HEK293 Expi suspension cultures using a three-plasmid transfection system. Viral particles were harvested from cell lysates and purified by affinity chromatography. Where indicated, full capsids were further enriched by iodixanol gradient ultracentrifugation. Vector genome yields at different production and purification stages were quantified by ddPCR. Data in panels a-d are shown as mean ± SD from 2 independent production batches. Biophysical analyses were performed on iodixanol-enriched full capsid preparations.

**a), b) Capsid population analysis by charge-detection mass spectrometry (CD-MS).** Particle populations were classified according to capsid mass as empty (black-3.5 – 4.0 Mda), partially filled (red – 4.00-4.51 MDa), full (green – 4.51-4.99 Mda) or overfilled (blue – 4.99-5.65 Mda). Presented are the data from 2 independent 2.8L production batches for **a)** AAV9 (AAV9.1; AAV9.2) and **b)** Bovine AAV (Bovine AAV1; Bovine AAV2).

Supplemental Figure 4. Pre-existing neutralizing antibodies against Bovine AAV, AAV2, AAV4, AAV5, and AAV9 in healthy adult cohorts from the USA versus Europe

Pre-existing neutralizing antibody responses against Bovine AAV, AAV2, AAV4, AAV5, and AAV9 were assessed in serum samples from healthy adults from the USA (*n* = 73) (**a,b,c**) and from Bern, Switzerland (*n* = 72) (**d**,**e,f**) using a cell-based transduction inhibition assay. Recombinant AAV vectors carrying an ss-CAG-GFP cassette were incubated with serially diluted human sera (1:20; 1:80; 1:320; 1:1,280; 1:5,120) for 1 h before addition to HEK-293T cells. Transduction efficiency was quantified 3 days later by flow cytometry. Normality was assessed using the Shapiro Wilk test. As the data were not normally distributed, differences in neutralizing antibody seroprevalence across AAV capsids were analyzed using the nonparametric Friedman test followed by Dunn multiple-comparisons test. Statistical significance: ∗*p*<0.05, ∗∗*p*<0.01, ∗∗∗*p*<0.001, ∗∗∗∗*p*<0.0001.

**a), d) Neutralization at the lowest serum dilution.** Percent neutralization at a serum dilution of 1:20, normalized to positive control wells (100%transduction) and negative control wells (0%transduction). Samples were classified as neutralizing if they inhibited transduction by at least 50 percent; the horizontal dashed line indicates this threshold. Each point represents one serum sample, shown as the mean of two technical replicates. Data are presented as median with interquartile range (IQR), 25th to 75th percentile: **a) USA cohort:** Bovine AAV (median: 39.55, IQR: 23.30 *to* 96.17; min-max: 2.78 *to* 100.0); AAV4 (median: 90.08, IQR: 79.69 *to* 95.14; min-max: 52.90 *to* 98.92); AAV2 (median: 98.85, IQR: 11.57 *to* 99.86; min-max:-1.11 *to* 99.98); AAV5 (median: 1.35, IQR:-0.28 *to* 5.85; min-max:-1.97 *to* 93.05); AAV9 (median:-14.10, IQR: 3.75 *to* 69.37; min-max:-8.83 *to* 99.92). **d) Switzerland cohort:** Bovine AAV (median: 10.73, IQR: 2.90 *to* 54.06; min-max:-3.84 *to* 99.9); AAV4 (median: 77.66, IQR: 63.78 *to* 85.48; min-max: 28.60 *to* 97.81); AAV2 (median: 33.18, IQR:-2.14 *to* 98.18; min-max:-4.81 *to* 99.85); AAV5 (median: 0.73, IQR:-1.33 *to* 6.84; min-max:-16.00 *to* 93.74); AAV9 (median:-1.26, IQR:-6.70 *to* 36.52; min-max:-11.36 *to* 99.98). Shown are only statistical differences relative to Bovine AAV. Statistical differences assessed by Friedman test followed by Dunn multiple-comparisons test: **a) US cohort**: Bovine AAV *vs* AAV4: *p*=0.528; Bovine AAV *vs* AAV2: *p*=0.185; Bovine *vs* AAV5: *p<*0.0001; Bovine AAV *vs* AAV9: *p*<0.001). **d) Switzerland cohort**: Bovine AAV *vs* AAV4: *p*=0.019; Bovine AAV *vs* AAV2: *p*>0.999; Bovine *vs* AAV5 / AAV9: *p<*0.0001)

**f) b), e) Seroprevalence of neutralizing antibodies across serum dilutions.** Heatmap showing the percentage of sera classified as neutralizing (≥ 50% inhibition of transduction) for each AAV serotype across the five serum dilutions tested in b) the USA and c) Switzerland cohorts.

**g) c), f) Neutralization across serum dilutions.** Percent neutralization of the indicated AAV capsids measured across the five serum dilutions for all sera samples tested in **c)** the USA (*n* = 73) and **g)** Switzerland (*n* = 72) cohorts. Each point represents one serum sample at a given dilution. The horizontal dashed line indicates the 50 percent neutralization threshold, with values above this line considered positive for neutralizing activity.

**f)**, h) Q Q plot analysis of neutralization data distribution at the lowest serum dilution (1:20) for **g) the US and h) the Swiss Bern cohorts.** Quantile quantile plots were used to compare the observed distribution of percent neutralization values for each AAV capsid with the theoretical normal distribution. Each point represents one serum sample, with symbols and colors indicating the corresponding capsid. The red dashed line indicates the expected distribution under normality. Systematic deviation of the data points from this line indicates non normality, which was confirmed by the Shapiro Wilk test for all capsids: **g) USA cohort** → Bovine AAV: *W*=0.84, *p*<0.0001; AAV4: *W*=0.87, *p*<0.0001; AAV2: *W*=0.67, *p*<0.0001; AAV5: *W*=0.43, *p*<0.0001; AAV9: *W*=0.81, *p*<0.0001. **h) Swiss Bern cohort** → Bovine AAV: *W*=0.74, *p*<0.0001; AAV4: *W*=0.94, *p*=0.0028; AAV2: *W*=0.73, *p*<0.0001; AAV5: *W*=0.62, *p*<0.0001; AAV9: *W*=0.71, *p*<0.0001.

Supplemental Figure 5. Implementation of a mouse model with established immunity to human **/ NHP-derived AAVs serotypes.**

**a) Study design.** Male C57BL/6j mice aged 9 to 11 weeks were injected intravenously with a pooled mixture of human and NHP-derived AAV empty capsids comprising AAV1, AAV2, AAV4, AAV5, AAV8, and AAV9. Two dose levels were tested as indicated in Supplemental Table 2 (*n* = 3 mice per group). An additional group receiving AAV formulation buffer only (No AAV), (*n* = 3) served as a negative control. Blood was collected from the lateral saphenous vein before injection and every 7 days thereafter for 49 days (marked with the gray arrows).

**b) Body weight monitoring**. Body weight was recorded 3 days before AAV administration and every 7 days throughout the 49-day study period. Data are shown as mean (SD) body weight in grams for n = 3 mice per group. Differences in body weight changes over time depending on the injected AAVs were analyzed using a two-way repeated-measures ANOVA with AAV treatment group as the between-subject factor and time as the within-subject factor, followed by Sidak multiple-comparisons test.

**c), d) Development of anti-AAV IgM and IgG titers.** Serum IgM (c) and IgG (d) responses against each AAV capsid present in the pool were measured at each blood collection timepoint by enzyme-linked immunosorbent assay (ELISA) using purified AAV capsids coated at 1.0 × 10^10^vp mL. Mouse sera were serially diluted in 11 three-fold steps from 1:50 to 1:2,952,450. Bound IgG or IgM antibodies were detected using anti-mouse HRP-conjugated secondary antibodies, followed by TMB substrate development. Absorbance was measured at 450 nm. Endpoint titers were defined as the highest serum dilution with an optical density, OD_450_, above the assay cutoff, calculated as two times the mean of 15 blank wells. Each dot represents one animal, and bars indicate mean (SD) for each group. The y axis shows the corresponding endpoint titer for each threefold dilution step. Shaded regions indicate low titers (1:50 to 1:450) (grey), intermediate titers (1:1,350 to 1:4,050) (orange) and high titers (1:12,150 to 1: 2,952,450) (red).

**d) Determination of appropriate MOIs for neutralization assay**. HEK-293T cells were transduced with AAV1, AAV2, AAV4, AAV5, AAV8, AAV9 or Bovine AAV vectors (carrying an ss-CAG-GFP cassette) using 11 two-fold serial dilutions ranging from 1.0 × 10^6^ to 9.77 × 10^2^ vg per cell. The percentage of GFP positive cells was quantified by flow cytometry 3 days later. MOIs yielding approximately 80 to 90 percent transduction and used for subsequent neutralization assays are highlighted by the grey box: AAV1 - 1.25 × 10^5^; AAV2 - 4.50 × 10^3^;AAV4 - 8.00 × 10^5^; AAV5 - 1.00 × 10^6^;AAV8 - 1.00 × 10^6^; and AAV9 - 8.00 × 10^5^vg per cell. Data are shown as mean (SD) of n = 2 technical replicates.

**e) Cell based neutralization assay.** Neutralizing activity against AAV1, AAV2, AAV4, AAV5, AAV8, and AAV9 was assessed using a serum-mediated transduction inhibition assay. For each experimental group, sera from all mice within the group were pooled and serially diluted (1:20; 1:80; 1:320; 1:1,280; 1:5,120) before pre incubation with recombinant AAV reporter vectors for 1 h at 37 °C. The serum AAV mixtures were then added to HEK-293T cells, and transduction efficiency was quantified 72 h later by flow cytometry as the percentage of GFP-positive cells. AAV vectors were used at MOIs previously selected to yield approximately 80 to 90 percent transduction in the absence of serum (panel e). Sera were classified as neutralizing if they reduced transduction by at least 50 percent at the lowest dilution tested (1:20).The dashed horizontal line indicates the 50 percent neutralization threshold. Points represent the mean transduction values obtained for each pooled serum group at the indicated dilutions. **Supplemental Figure 6. Evaluation of humoral immunity induced by AAV pool administration and cross neutralization of Bovine AAV.**

**a) Development of anti-AAV IgM and b) IgG responses after intravenous administration of the AAV pool.** Serum IgM **(a)** & IgG **(b)** titers against each AAV capsid present in the pool were measured at each blood collection timepoint by ELISA. Plates were coated with purified AAV capsids at 1.0 × 10^10^vp mL and incubated with mouse sera serially diluted in 11 three-fold steps, from 1:50 to 1:2,952,450. Bound antibodies were detected using HRP conjugated anti mouse secondary antibodies and TMB substrate, and absorbance was measured at 450 nm. Endpoint titers were defined as the highest serum dilution with an *OD*_450_ above the assay cutoff, calculated as two times the mean of 15 blank replicates. Plates coated with buffer (No coat) served as negative control plates. Each dot represents one animal, and bars indicate mean (SD) for the group (n=5 mice/ group). The y axis shows the corresponding endpoint titer for each threefold dilution step. Shaded regions indicate low titers (1:50 to 1:450), intermediate titers (1:1,350 to 1:4,050) and high titers (1:12,150 to 1: 2,952,450).

**b) Neutralization of Bovine AAV by sera from mice immunized with the AAV pool.** Neutralizing activity against Bovine AAV was assessed by cell based transduction inhibition assay. Sera were collected at the experimental endpoint (*day* 49) from mice that received the AAV pool alone, No AAV (negative control), or the AAV pool (Day 0) followed by Bovine AAV (Day 21) (positive control). For each experimental group, sera collected at day 49 post immunization were serially diluted starting at 1:20 before incubation with Bovine AAV at an MOI of 1.0 × 10^5^vg per cell for 1 h at 37 °C. Transduction efficiency was quantified 72 h later by flow cytometry as the percentage of GFP positive cells.

**c) Neutralization of AAV4 and Bovine AAV by sera from AAV4 immunized mice.** Male C57BL/6j mice (*n* = 3 per group) were injected intravenously with AAV4 at either 5.0 × 10^11^vg kg, corresponding to 1.25 × 10^10^vg per 25 g mouse, or 5.0 × 10^12^vg kg, corresponding to 1.25 × 10^11^vg per 25 g mouse. Twenty-one days later, mice were euthanized and blood was collected by intracardiac puncture. Neutralizing activity against AAV4 and Bovine AAV was assessed in mouse sera collected at day 28 post immunization, using the cell-based transduction inhibition assay described above (*Supplementary Fig*. 6*c*).

## Tables

**Table 1.** Healthy Donor demographics.

| Cohort Origin | Nr samples | Sex donors | Age range |  | Ethnicity |  |  |  | Health status |
| --- | --- | --- | --- | --- | --- | --- | --- | --- | --- |
|  |  |  | 20-60 years | 61-75 years | Black | White | Asian | Other race |  |
| USA | 73 | 31 females | 28 | 3 | 4 | 22 | 2 | 3 | healthy |
|  |  | 42 males | 34 | 8 | 7 | 18 | 8 | 9 |  |
| Europe (Switzerland, Bern) | 72 | 20 females | 16 | 4 | N/A |  |  |  | healthy |
|  |  | 36 males | 27 | 9 |  |  |  |  |  |
|  |  | 16 N/A | adults |  |  |  |  |  |  |

## Supplemental Tables (Methods)

**Supplemental Table 3.** Rep-Cap information for all AAV serotypes produced.

| AAV serotype | Rep-Cap | Size (bp) |
| --- | --- | --- |
| AAV1 | Rep2Cap1 | 7279 |
| AAV2 | Rep2Cap2 | 7276 |
| AAV4 | Rep2Cap4 | 7273 |
| AAV5 | Rep2Cap5 | 7243 |
| AAV8 | Rep2Cap8 | 7432 |
| AAV9 | Rep2Cap9 | 7279 |
| AAV2-ESGHGYF | Rep2Cap2-ESGHGYF | 7303 |
| Bovine AAV | Rep2CapBovine AAV | 7279 |

**Supplemental Table 4.** Density of different cell types cultured in 96 or 384 well plate at plating day (generally 24h before AAV transduction)

| Cell type | Cell density at plating (96 well) | Cell density at plating (384 well) |
| --- | --- | --- |
| HEK-293T | 15 000 cells/ well | 3 000 cells/ well |
| Small airway epithelial cells | 7 000 cells / well | 1 700 cells/ well |
| Bronchial / tracheal epithelial cells | 7 000 cells / well | 1 700 cells/ well |
| Pulmonary alveolar epithelial cells | 20 000 cells / well | 4000 cells/ well |
| Lung microvascular endothelial cells | 5000 cells/ well | 1000 cells / well |

