## Supplemental Figures & Tables for "Bovine AAV - a promising vector for pulmonary gene therapy"

**Figure S1**

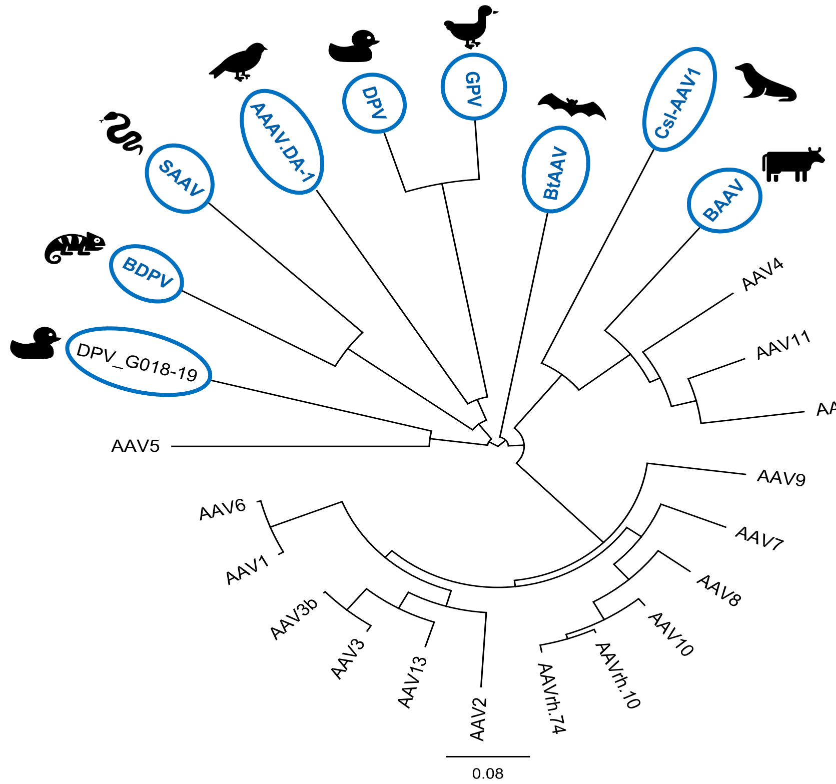


**Table S1. Table displaying VP1 amino acid sequence similarity** (shown in percentages) between phylogenetically distant and human / non-human primate-derived AAVs. Green values indicate low similarity, yellow intermediate and orange/red high similarity.

| % AAV identity at VP1 level | AAAV.DA-1 | AAV1 | AAV2 | AAV3 | AAV3b | AAV4 | AAV5 | AAV6 | AAV7 | AAV8 | AAV9 | AAV10 | AAV11 | AAV12 | AAV13 | AAVrh.10 | AAVrh.74 | BAAV | BDPV | BtAAV | Csl-AAV1 | DPV | DPV_G018-19 | GPV | SAAV |
|---|---|---|---|---|---|---|---|---|---|---|---|---|---|---|---|---|---|---|---|---|---|---|---|---|---|
| **AAAV.DA-1** | | 55.6 | 56.21 | 57.47 | 57.73 | 52.37 | 53.29 | 55.87 | 57.31 | 56.12 | 56.19 | 56.65 | 53.23 | 50.97 | 57.09 | 56.52 | 56.38 | 52.23 | 50.99 | 52.76 | 50.13 | 55.15 | 58.13 | 56.09 | 48.03 |
| **AAV1** | 55.6 | | 83.29 | 86.43 | 86.84 | 63.05 | 57.14 | 99.18 | 84.96 | 83.88 | 82.23 | 84.69 | 66.49 | 59.89 | 86.68 | 84.82 | 84.01 | 59.95 | 54.41 | 59.81 | 55.45 | 52.98 | 58.01 | 52.85 | 52.46 |
| **AAV2** | 56.21 | 83.29 | | 87.23 | 87.77 | 59.92 | 57.14 | 83.42 | 82.38 | 82.79 | 81.68 | 83.74 | 62.55 | 59.05 | 88.03 | 84.01 | 83.88 | 58.16 | 53.48 | 59.49 | 53.13 | 52.39 | 57.09 | 52.25 | 51 |
| **AAV3** | 57.47 | 86.43 | 87.23 | | 99.18 | 61.98 | 57.2 | 86.57 | 84.28 | 85.12 | 83.47 | 85.39 | 63.67 | 59.63 | 93.61 | 85.12 | 84.71 | 59.41 | 54.21 | 59.41 | 54.52 | 53.64 | 57.94 | 53.64 | 51.99 |
| **AAV3b** | 57.73 | 86.84 | 87.77 | 99.18 | | 62.52 | 57.47 | 86.97 | 84.82 | 85.52 | 83.88 | 85.79 | 64.08 | 60.16 | 93.89 | 85.52 | 85.12 | 59.95 | 54.61 | 59.81 | 54.79 | 53.91 | 58.33 | 53.91 | 52.13 |
| **AAV4** | 52.37 | 63.05 | 59.92 | 61.98 | 62.52 | | 51.4 | 62.92 | 62.75 | 62.88 | 61.9 | 62.88 | 81.36 | 78.61 | 63.62 | 62.88 | 62.75 | 76.66 | 48.75 | 55.91 | 59.66 | 50.2 | 51.96 | 50.2 | 47.76 |
| **AAV5** | 53.29 | 57.14 | 57.14 | 57.2 | 57.47 | 51.4 | | 57.28 | 56.99 | 56.85 | 56.47 | 56.32 | 51.94 | 50.66 | 57.49 | 56.59 | 56.18 | 53.06 | 49.53 | 54.16 | 48.21 | 51.55 | 61.37 | 52.09 | 47 |
| **AAV6** | 55.87 | 99.18 | 83.42 | 86.57 | 86.97 | 62.92 | 57.28 | | 85.09 | 84.01 | 82.09 | 84.82 | 66.35 | 59.76 | 87.09 | 84.96 | 84.15 | 59.81 | 54.41 | 59.81 | 55.32 | 52.72 | 58.28 | 52.72 | 52.33 |
| **AAV7** | 57.31 | 84.96 | 82.38 | 84.28 | 84.82 | 62.75 | 56.99 | 85.09 | | 87.96 | 81.73 | 88.36 | 66.31 | 60.26 | 84.82 | 88.63 | 88.23 | 59.65 | 54 | 61.07 | 54.77 | 54.03 | 57.07 | 54.16 | 51.66 |
| **AAV8** | 56.12 | 83.88 | 82.79 | 85.12 | 85.52 | 62.88 | 56.85 | 84.01 | 87.96 | | 85.23 | 93.36 | 65.24 | 60.26 | 85.37 | 93.5 | 93.36 | 58.32 | 54 | 59.6 | 55.44 | 54.56 | 57.07 | 54.69 | 51.93 |
| **AAV9** | 56.19 | 82.23 | 81.68 | 83.47 | 83.88 | 61.9 | 56.47 | 82.09 | 81.73 | 85.23 | | 85.64 | 62.78 | 59.42 | 83.85 | 85.5 | 85.5 | 58 | 53.21 | 59.41 | 54.52 | 53.25 | 56.95 | 53.25 | 51.93 |
| **AAV10** | 56.65 | 84.69 | 83.74 | 85.39 | 85.79 | 62.88 | 56.32 | 84.82 | 88.36 | 93.36 | 85.64 | | 65.91 | 60.13 | 86.18 | 98.37 | 97.83 | 59.39 | 54.8 | 60.4 | 55.17 | 54.29 | 57.2 | 54.03 | 52.46 |
| **AAV11** | 53.23 | 66.49 | 62.55 | 63.67 | 64.08 | 81.36 | 51.94 | 66.35 | 66.31 | 65.24 | 62.78 | 65.91 | | 83.4 | 64.03 | 65.64 | 65.11 | 78.97 | 50.33 | 55.38 | 59.34 | 50.33 | 52.03 | 49.67 | 47.11 |
| **AAV12** | 50.97 | 59.89 | 59.05 | 59.63 | 60.16 | 78.61 | 50.66 | 59.76 | 60.26 | 60.26 | 59.42 | 60.13 | 83.4 | | 59.84 | 60 | 59.74 | 74.53 | 48.89 | 52.31 | 57.89 | 49.03 | 49.81 | 49.03 | 46.61 |
| **AAV13** | 57.09 | 86.68 | 88.03 | 93.61 | 93.89 | 63.62 | 57.49 | 87.09 | 84.82 | 85.37 | 83.85 | 86.18 | 64.03 | 59.84 | | 86.04 | 85.64 | 60.29 | 54.42 | 59.84 | 55.13 | 53.78 | 58.62 | 53.52 | 51.94 |
| **AAVrh.10** | 56.52 | 84.82 | 84.01 | 85.12 | 85.52 | 62.88 | 56.59 | 84.96 | 88.63 | 93.5 | 85.5 | 98.37 | 65.64 | 60 | 86.04 | | 98.92 | 58.99 | 54.4 | 60.4 | 55.17 | 54.29 | 57.2 | 54.03 | 52.46 |
| **AAVrh.74** | 56.38 | 84.01 | 83.88 | 84.71 | 85.12 | 62.75 | 56.18 | 84.15 | 88.23 | 93.36 | 85.5 | 97.83 | 65.11 | 59.74 | 85.64 | 98.92 | | 59.12 | 54.53 | 60.13 | 55.04 | 54.43 | 56.54 | 54.29 | 52.72 |
| **BAAV** | 52.23 | 59.95 | 58.16 | 59.41 | 59.95 | 76.66 | 53.06 | 59.81 | 59.65 | 58.32 | 58 | 59.39 | 78.97 | 74.53 | 60.29 | 58.99 | 59.12 | | 49.08 | 57.45 | 59.76 | 50 | 50.91 | 50.26 | 47.31 |
| **BDPV** | 50.99 | 54.41 | 53.48 | 54.21 | 54.61 | 48.75 | 49.53 | 54.41 | 54 | 54 | 53.21 | 54.8 | 50.33 | 48.89 | 54.42 | 54.4 | 54.53 | 49.08 | | 51.99 | 46.9 | 49.21 | 50.4 | 49.61 | 68.63 |
| **BtAAV** | 52.76 | 59.81 | 59.49 | 59.41 | 59.81 | 55.91 | 54.16 | 59.81 | 61.07 | 59.6 | 59.41 | 60.4 | 55.38 | 52.31 | 59.84 | 60.4 | 60.13 | 57.45 | 51.99 | | 52.18 | 51.72 | 52.64 | 52.25 | 51.06 |
| **Csl-AAV1** | 50.13 | 55.45 | 53.13 | 54.52 | 54.79 | 59.66 | 48.21 | 55.32 | 54.77 | 55.44 | 54.52 | 55.17 | 59.34 | 57.89 | 55.13 | 55.17 | 55.04 | 59.76 | 46.9 | 52.18 | | 48.21 | 48.89 | 48.47 | 45.59 |
| **DPV** | 55.15 | 52.98 | 52.39 | 53.64 | 53.91 | 50.2 | 51.55 | 52.72 | 54.03 | 54.56 | 53.25 | 54.29 | 50.33 | 49.03 | 53.78 | 54.29 | 54.43 | 50 | 49.21 | 51.72 | 48.21 | | 52.07 | 87.57 | 48.75 |
| **DPV_G018-19** | 58.13 | 58.01 | 57.09 | 57.94 | 58.33 | 51.96 | 61.37 | 58.28 | 57.07 | 57.07 | 56.95 | 57.2 | 52.03 | 49.81 | 58.62 | 57.2 | 56.54 | 50.91 | 50.4 | 52.64 | 48.89 | 52.07 | | 53.14 | 48.03 |
| **GPV** | 56.09 | 52.85 | 52.25 | 53.64 | 53.91 | 50.2 | 52.09 | 52.72 | 54.16 | 54.69 | 53.25 | 54.03 | 49.67 | 49.03 | 53.52 | 54.03 | 54.29 | 50.26 | 49.61 | 52.25 | 48.47 | 87.57 | 53.14 | | 49.41 |
| **SAAV** | 48.03 | 52.46 | 51 | 51.99 | 52.13 | 47.76 | 47 | 52.33 | 51.66 | 51.93 | 51.93 | 52.46 | 47.11 | 46.61 | 51.94 | 52.46 | 52.72 | 47.31 | 68.63 | 51.06 | 45.59 | 48.75 | 48.03 | 49.41 | |

| % |
|---|
| 100 |
| 90 |
| 80 |
| 70 |
| 60 |
| 50 |
| 40 |

**Figure S2**

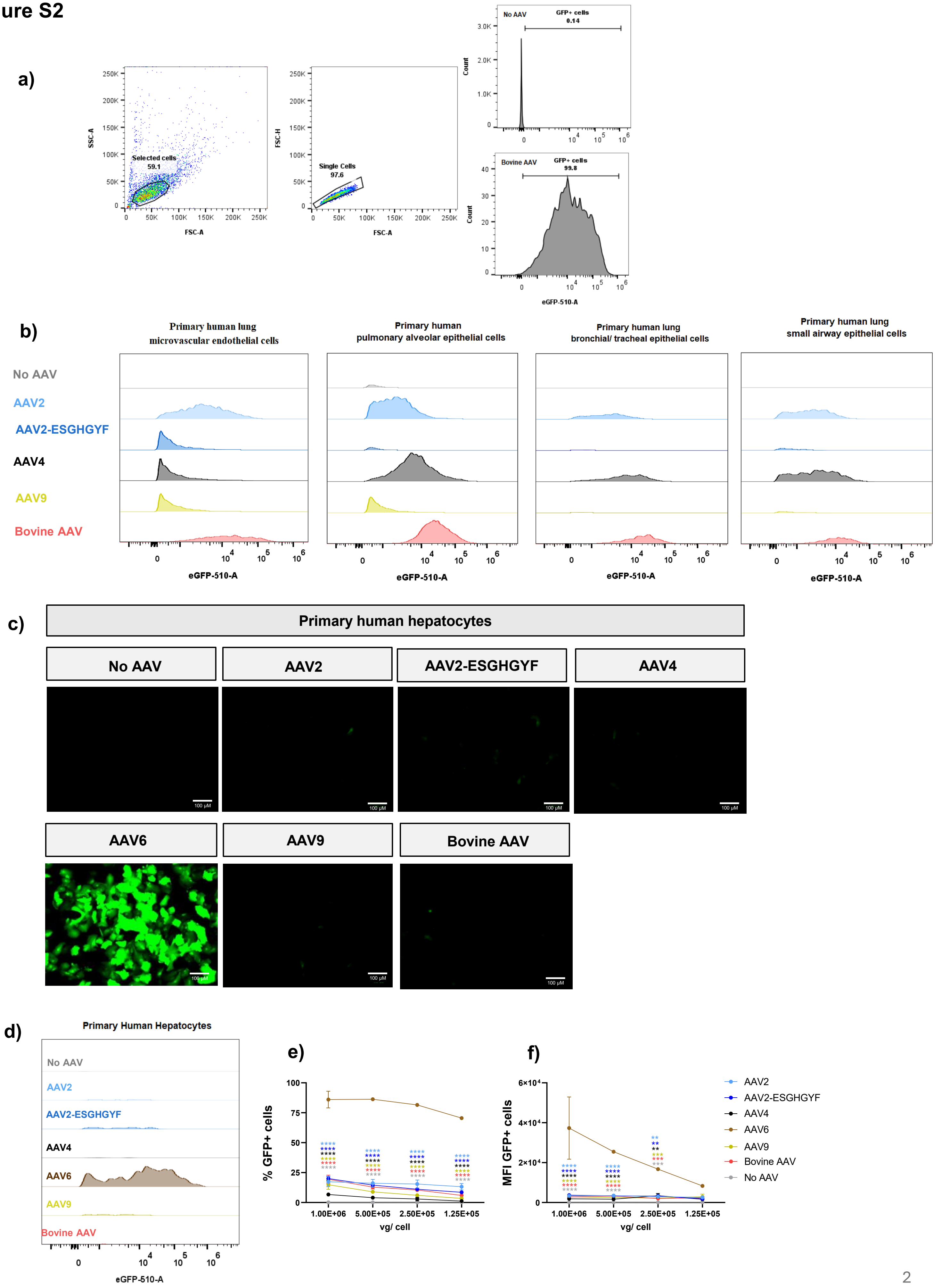
a)
SSC-A
FSC-A
Selected cells
59.1
FSC-H
Single Cells
97.6
No AAV
GFP+ cells
0.14
Bovine AAV
GFP+ cells
99.8
Count
eGFP-510-A
b)
Primary human lung microvascular endothelial cells
Primary human pulmonary alveolar epithelial cells
Primary human lung bronchial/ tracheal epithelial cells
Primary human lung small airway epithelial cells
No AAV
AAV2
AAV2-ESGHGYF
AAV4
AAV9
Bovine AAV
eGFP-510-A
c)
Primary human hepatocytes
No AAV
AAV2
AAV2-ESGHGYF
AAV4
AAV6
AAV9
Bovine AAV
100 µM
d)
Primary Human Hepatocytes
No AAV
AAV2
AAV2-ESGHGYF
AAV4
AAV6
AAV9
Bovine AAV
eGFP-510-A
e)
% GFP+ cells
vg/ cell
1.00E+06
5.00E+05
2.50E+05
1.25E+05
f)
MFI GFP+ cells
vg/ cell
AAV2
AAV2-ESGHGYF
AAV4
AAV6
AAV9
Bovine AAV
No AAV

### Figure S3

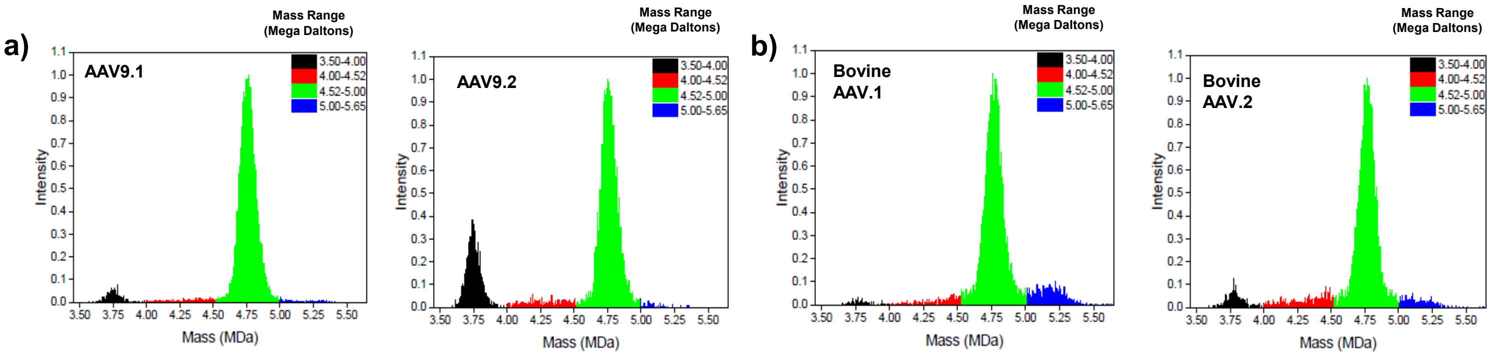


#### Table S2 - Biophysical characteristics of AAV9 and Bovine AAV

| | Sample Information | | SEC-MALS | | | CD-MS | | | | HT CE-SDS | | | | | Capsid Unfolding | | Genome Release | | |
|---|---|---|---|---|---|---|---|---|---|---|---|---|---|---|---|---|---|---|---|
| Experiment N° | AAV Serotype | Cargo Size | HMW [%] | Full [%] | Particle Conc. [VP/mL] | Empty [%] | Full [%] | Over-filled [%] | Capsid [%] | LMWs [%] | VP composition | | | VP3:VP2:VP1 ratio | ON [°C] | Tm1 [°C] | ON [°C] | DNA Tm1 [°C] | DNA Tm2 [°C] |
| | | | | | | | | | | | VP3 | VP2 | VP1 | | | | | | |
| 1 | AAV9.1 | 2983 | 1.0 | 78 | 2.6E+13 | 44.7 | 54.8 | 0.5 | 78 | 3.3 | 49.6 | 5.2 | 5.2 | 9.5: 1: 1 | 71 | 76 | 49 | 62 | 72 |
| 2 | AAV9.2 | | 3.3 | 68 | 4.0E+13 | 20.6 | 73.8 | 0.9 | 94.8 | 7.4 | 50.0 | 5.7 | 4.3 | 12: 1: 1 | 74 | 78 | 52 | 62 | 72 |
| 1 | Bovine AAV.1 | 2983 | 0.8 | 80 | 2.8E+13 | 25.2 | 74.4 | 0.4 | 80 | 6.7 | 49.8 | 5.4 | 4.8 | 9.3: 1: 0.9 | 64 | 70 | 40 | 52 | 65 |
| 2 | Bovine AAV.2 | | 1.2 | 86 | 3.7E+13 | 5.0 | 82.3 | 4.8 | 87.8 | 8.3 | 50.2 | 5.7 | 4.1 | 12: 1: 1 | 64 | 70 | 42 | 52 | 65 |

HMW = high molecular weight; LMW = low molecular weight; SEC-MALS = Size-exclusion chromatography with multi-angle light scattering detection; HT CE-SDS = High-throughput microfluidic chip-based capillary gel electrophoresis; DSF = Differential Scanning Fluorimetry; ON = onset temperature; Tm = melting temperature; VP = viral particle

### Figure S4

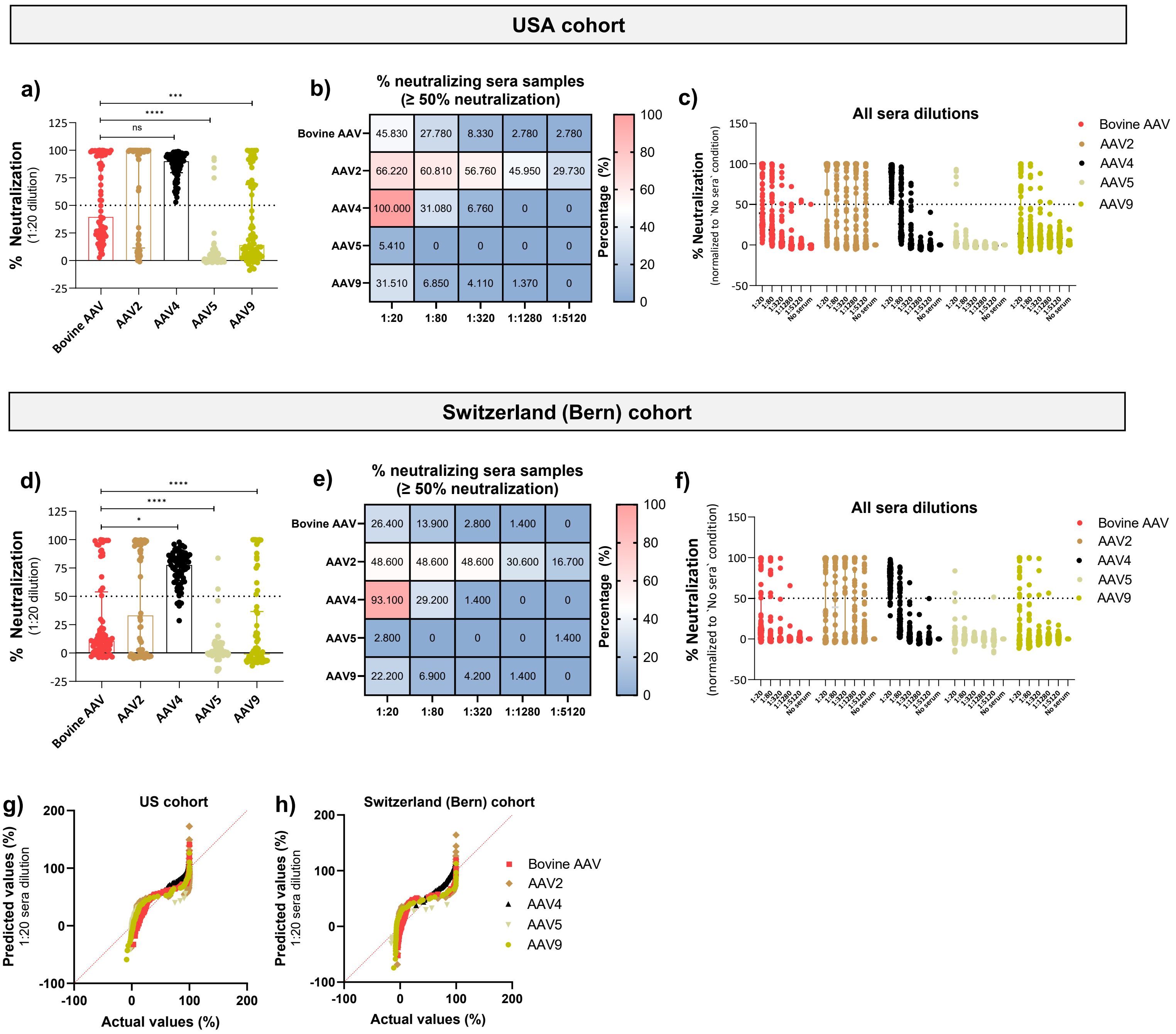
USA cohort
a)
% Neutralization
(1:20 dilution)
125
100
75
50
25
0
-25
***
****
ns
Bovine AAV
AAV2
AAV4
AAV5
AAV9
b)
% neutralizing sera samples
(≥ 50% neutralization)
Bovine AAV 45.830 27.780 8.330 2.780 2.780
AAV2 66.220 60.810 56.760 45.950 29.730
AAV4 100.000 31.080 6.760 0 0
AAV5 5.410 0 0 0 0
AAV9 31.510 6.850 4.110 1.370 0
1:20 1:80 1:320 1:1280 1:5120
Percentage (%)
100
80
60
40
20
0
c)
All sera dilutions
% Neutralization
(normalized to `No sera` condition)
150
100
50
0
-50
1:20
1:80
1:320
1:1280
1:5120
No serum
Bovine AAV
AAV2
AAV4
AAV5
AAV9
Switzerland (Bern) cohort
d)
% Neutralization
(1:20 dilution)
125
100
75
50
25
0
-25
****
****
*
Bovine AAV
AAV2
AAV4
AAV5
AAV9
e)
% neutralizing sera samples
(≥ 50% neutralization)
Bovine AAV 26.400 13.900 2.800 1.400 0
AAV2 48.600 48.600 48.600 30.600 16.700
AAV4 93.100 29.200 1.400 0 0
AAV5 2.800 0 0 0 1.400
AAV9 22.200 6.900 4.200 1.400 0
1:20 1:80 1:320 1:1280 1:5120
Percentage (%)
100
80
60
40
20
0
f)
All sera dilutions
% Neutralization
(normalized to `No sera` condition)
150
100
50
0
-50
1:20
1:80
1:320
1:1280
1:5120
No serum
Bovine AAV
AAV2
AAV4
AAV5
AAV9
g)
US cohort
Predicted values (%)
1:20 sera dilution
200
100
0
-100
-100
0
100
200
Actual values (%)
h)
Switzerland (Bern) cohort
Predicted values (%)
1:20 sera dilution
200
100
0
-100
-100
0
100
200
Actual values (%)
Bovine AAV
AAV2
AAV4
AAV5
AAV9

### Figure S5

a)

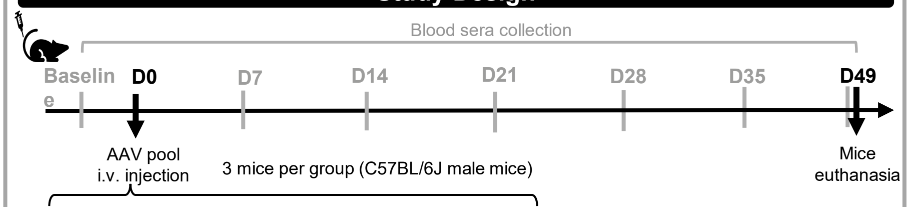


| AAV Serotype (no cargo) | No AAV Ctrl | Low Dose (vp/kg) | Intermediate Dose (vp/kg) |
|---|---|---|---|
| AAV1 | 0 | 5.00E+11 | 8.00E+11 |
| AAV2 | 0 | 2.50E+10 | 2.50E+11 |
| AAV4 | 0 | 2.00E+11 | 5.00E+11 |
| AAV5 | 0 | 2.50E+11 | 5.00E+11 |
| AAV8 | 0 | 2.00E+11 | 8.00E+11 |
| AAV9 | 0 | 5.00E+10 | 5.00E+11 |
| **Total AAV pool (vp/kg)** | **0** | **1.03E+12** | **3.35E+12** |

b)

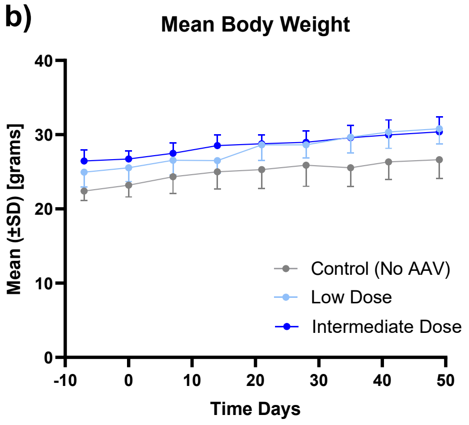


c)

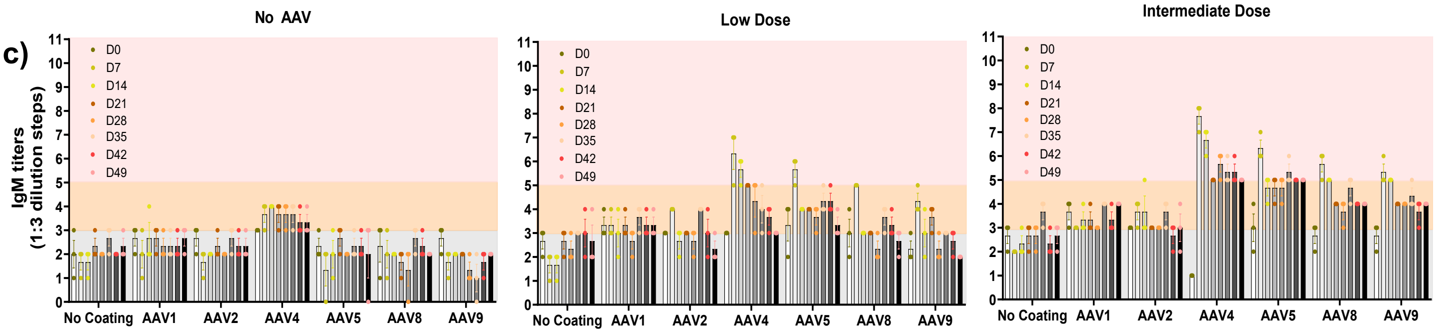


| Sera dilution | Dilution step |
|---|---|
| 1 : 50 | 1 |
| 1 : 150 | 2 |
| 1 : 450 | 3 |
| 1 : 1350 | 4 |
| 1: 4050 | 5 |
| 1 : 12150 | 6 |
| 1 : 36450 | 7 |
| 1 : 109350 | 8 |
| 1 : 328050 | 9 |
| 1 : 984150 | 10 |
| 1 : 2952450 | 11 |

d)

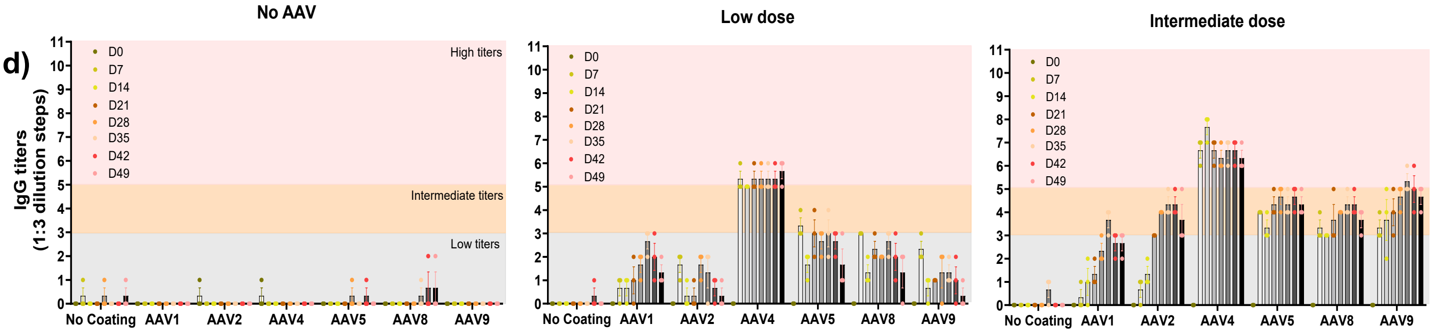


e)

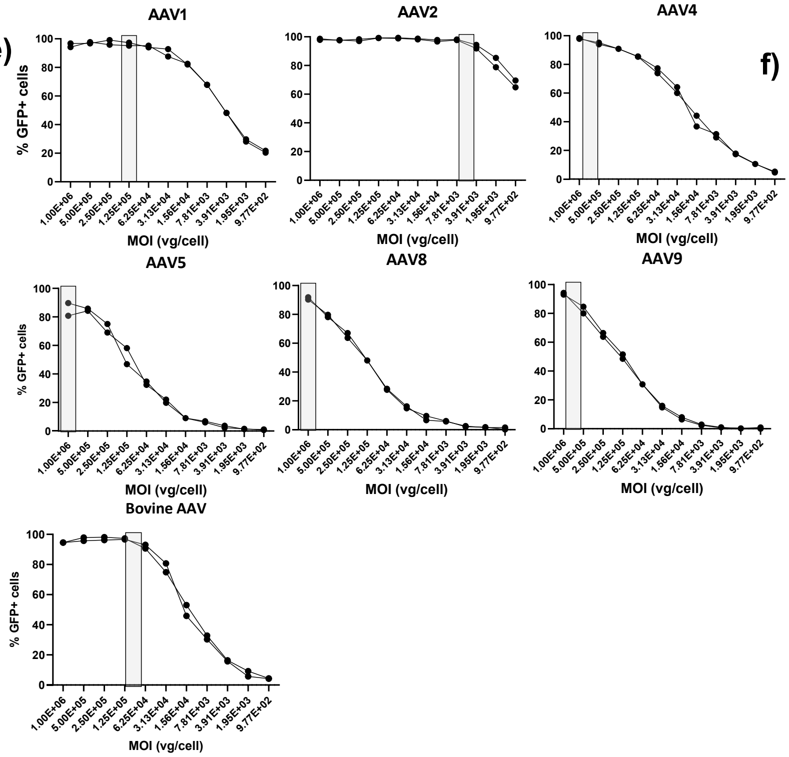


f)

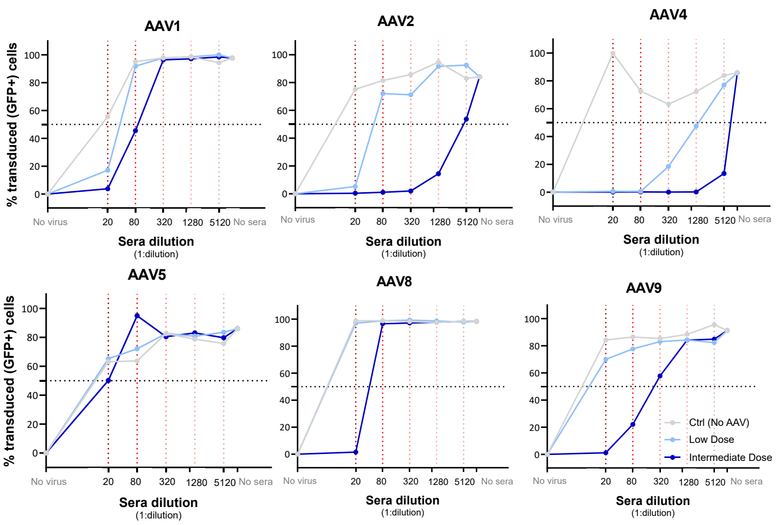

### Figure S6

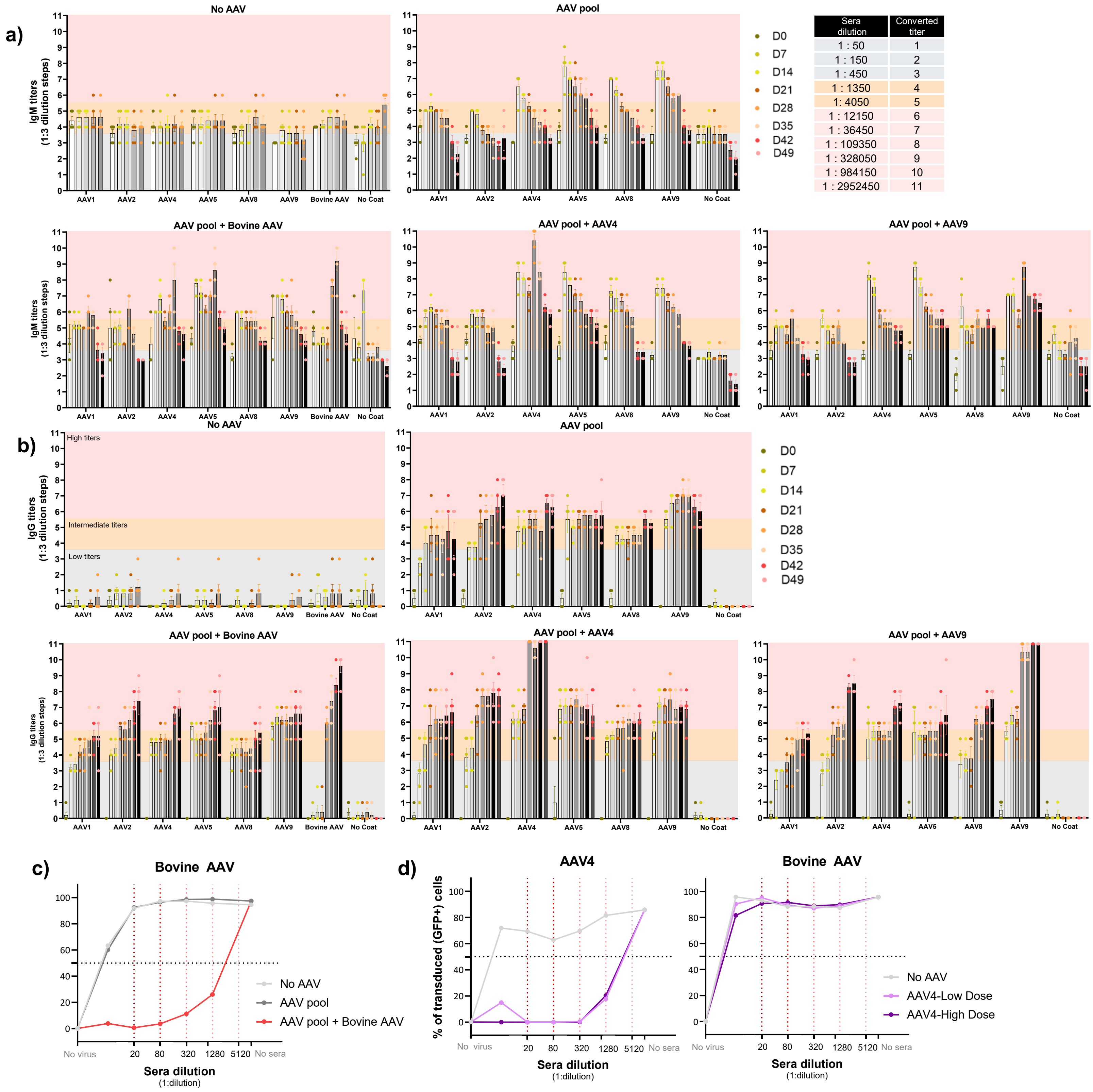


| Sera dilution | Converted titer |
|---|---|
| 1 : 50 | 1 |
| 1 : 150 | 2 |
| 1 : 450 | 3 |
| 1 : 1350 | 4 |
| 1: 4050 | 5 |
| 1 : 12150 | 6 |
| 1 : 36450 | 7 |
| 1 : 109350 | 8 |
| 1 : 328050 | 9 |
| 1 : 984150 | 10 |
| 1 : 2952450 | 11 |
